# KIF1C, an RNA transporting kinesin-3, undergoes liquid-liquid phase separation through its C-terminal disordered domain

**DOI:** 10.1101/2023.10.23.563538

**Authors:** Qi Geng, Jakia Jannat Keya, Takashi Hotta, Kristen J Verhey

## Abstract

The spatial distribution of mRNA is critical for local control of protein production. Recent studies have identified the kinesin-3 family member KIF1C as an RNA transporter. However, it is not clear how KIF1C interacts with RNA molecules. Here, we show that KIF1C’s C-terminal tail domain is an intrinsically disordered region (IDR) containing a prion-like domain (PLD) that is unique compared to the C-terminal tails of other kinesin family members. In cells, KIF1C constructs undergo reversible formation of dynamic puncta that display physical properties of liquid condensates and incorporate RNA molecules in a sequence-selective manner. The IDR is necessary and sufficient for driving liquid-liquid phase separation (LLPS) but the condensate properties can be modulated by adjacent coiled-coil segments. The purified KIF1C IDR domain undergoes LLPS *in vitro* at near-endogenous nM concentrations in a salt-dependent manner. Deletion of the IDR abolished the ability of KIF1C to undergo LLPS and disrupted the distribution of mRNA cargoes to the cell periphery. Our work thus uncovers an intrinsic correlation between the LLPS activity of KIF1C and its role as an RNA transporter. In addition, as the first kinesin motor reported to undergo LLPS, our work reveals a previously uncharacterized mode of motor-cargo interaction that extends our understanding of the behavior of cytoskeletal motor proteins.

## Introduction

Subcellular targeting of mRNAs is prevalent and can be used to spatially and temporally regulate the processing, translation, and stability of mRNA in cells^1–5^. For example, β-actin mRNA is distributed to the leading edge of migrating cells where local translation can produce abundant actin required for lamellipodium formation^6–11^. Subsequent studies found numerous mRNAs that are enriched in protrusions of migratory cells^12–14^, such as the small GTPase Rab13 whose mRNA distribution affects its co-translational association with different protein complexes during cell migration and tissue morphogenesis^15–17^. Subcellular targeting of mRNAs has also been shown to be critical in differentiated cells such as oligodendrocytes and neurons^18–21^.

Subcellular targeting of mRNAs has several advantages over localization of their protein products, such as the ability to regulate translation in response to local stimuli, and to prevent proteins from undesired interactions or premature functions before reaching their destination.

In general, there are two mechanisms for subcellular targeting or compartmentalization of mRNAs. First, mRNAs can be enriched into membrane-less bodies called RNA granules through liquid-liquid phase separation (LLPS)^22–24^. LLPS is the process of molecules working against entropy to cluster into a dense phase termed a biomolecular condensate that is separated from the surrounding diffusive phase. LLPS is driven by weak, multivalent interactions between molecules that typically contain intrinsically disordered regions with low complexity sequences^25,26^. Their highly dynamic nature and sensitivity to the chemical environment empowers LLPS condensates to regulate translational activity as well as the degradation of mRNAs^25–28^.

Second, mRNAs can be targeted via active transport by molecular motor proteins^29–33^. For microtubule-mediated long-distance transport, <u>ki</u>nesin family (KIF) proteins distribute cargoes towards microtubule plus ends in the cell periphery, while dynein proteins drive transport in the opposite direction. mRNAs are thought to form complexes with molecular motors via RNA-binding adaptor proteins such as Egalitarian^34^. During transport, mRNAs can present as single molecules or in RNA granules^35–40^. mRNAs can also be transported by hitchhiking on organelles such as endosomes and lysosomes^35,41,42^. How motor proteins bind to and transport specific mRNAs and other RNA species remains an important question in the field.

Recent work has shown that KIF1C, a ubiquitously expressed member of the kinesin-3 family, functions in mRNA transport in numerous cell types^43–46^. KIF1C had previously been implicated in the transport of Golgi-derived vesicles, integrins, and secretory vesicles^47–49^, and thus appears to be a kinesin protein capable of transporting both membrane-bound organelles and membrane- less complexes.

During cell migration, specific mRNAs are localized to protrusions at the leading edge^13,50^. While the mRNAs of most motor proteins localize randomly throughout the cytoplasm, the mRNAs of three kinesins were found to localize specifically to protrusions: the kinesin-1 *KIF5B*, the kinesin-3 *KIF1C*, and the kinesin-4 *KIF4A*^12^. Interestingly, KIF1C was the only kinesin where the protein and mRNA co-localized at the protrusion^12^. Immunoprecipitation of KIF1C identified a number of co-precipitated mRNAs including the protrusion-localized mRNAs of *NET1*, *TRAK2*, *RAB13*, and *KIF1C* itself^43^. Two-color imaging showed that KIF1C directly transports mRNAs and drives the formation of peripheral, multimeric RNA clusters^43^. Recently, KIF1C was found to interact with several members of the exon junction complex (EJC), which binds to mRNAs throughout their life cycle, and loss of KIF1C activity resulted in decreased transport of EJC components along neuronal processes^44^. KIF1C was also found to interact with Muscleblind-like (MBNL) proteins involved in splicing, polyadenylation and localization of mRNAs in numerous cell types^46^. Whether KIF1C binds to adaptor proteins to recruit specific mRNAs or binds to specific mRNAs that are associated with adaptor proteins is largely unknown. Support for the former possibility comes from recent finding that KIF1C’s association with MBNL occurs in an RNA-independent manner^46^. However, several studies support the latter possibility as mRNA–protein cross-linking approaches identified KIF1C as a direct binder of mRNAs^44,51,52^ and binding between KIF1C and EJC components is RNA-mediated^44^.

In this study, we show that the C-terminal tail domain of KIF1C is an intrinsically disordered region (IDR) with low-complexity sequence and a putative prion-like domain (PLD). The IDR is necessary and sufficient for LLPS and the formation of KIF1C condensates with liquid or hydrogel properties. To our knowledge, this is the first kinesin known to undergo LLPS. Synthesized RNA oligos and endogenous *Rab13* mRNA can be enriched into KIF1C condensates in a sequence-selective manner. Overall, our findings provide a novel mechanism of motor-RNA interaction through LLPS, highlighting the behavior of the disordered KIF1C C-terminal tail domain that is unique among the kinesin superfamily.

## Results

### KIF1C forms dynamic puncta via its C-terminal tail domain

To probe the localization of KIF1C in mammalian cells, we expressed fluorescently-tagged full-length KIF1C (Fig. 1 A) in COS-7 cells by transient transfection. KIF1C localized diffusely throughout the cell as well as to punctate structures (Fig. 1 B). Other fluorescently-tagged kinesin-3 family members showed similar localization patterns as expected: KIF1Bβ localized diffusely and to lysosomes ^53^, KIF13B localized diffusely and to small secretory vesicles ^54^, and KIF16B localized diffusely and to endosomes ^55^ (Fig. 1 B). Interestingly, the KIF1C puncta appeared to be heterogenous in size whereas the other kinesin-3 puncta appeared more homogeneous in size. Indeed, quantification showed that the KIF1C puncta ranged in size from 0.30 to 6.13 μm in diameter (average 1.31 ± 0.91 μm, Fig. 1 C) whereas KIF16B puncta had a more uniform size distribution (ranging from 0.35 to 1.48 μm in diameter, average 0.75 ± 0.17 μm, Fig. 1 C). We wondered whether the larger puncta could be formed by aggregation or fusion of smaller puncta. By live-cell imaging, KIF1C puncta were observed to undergo fission into smaller puncta which could then undergo fusion events to generate larger puncta over a time scale of several seconds (Fig. 1 D; Video 1, Video 2). This suggests that the KIF1C puncta are dynamic in nature.

**Figure 1.**
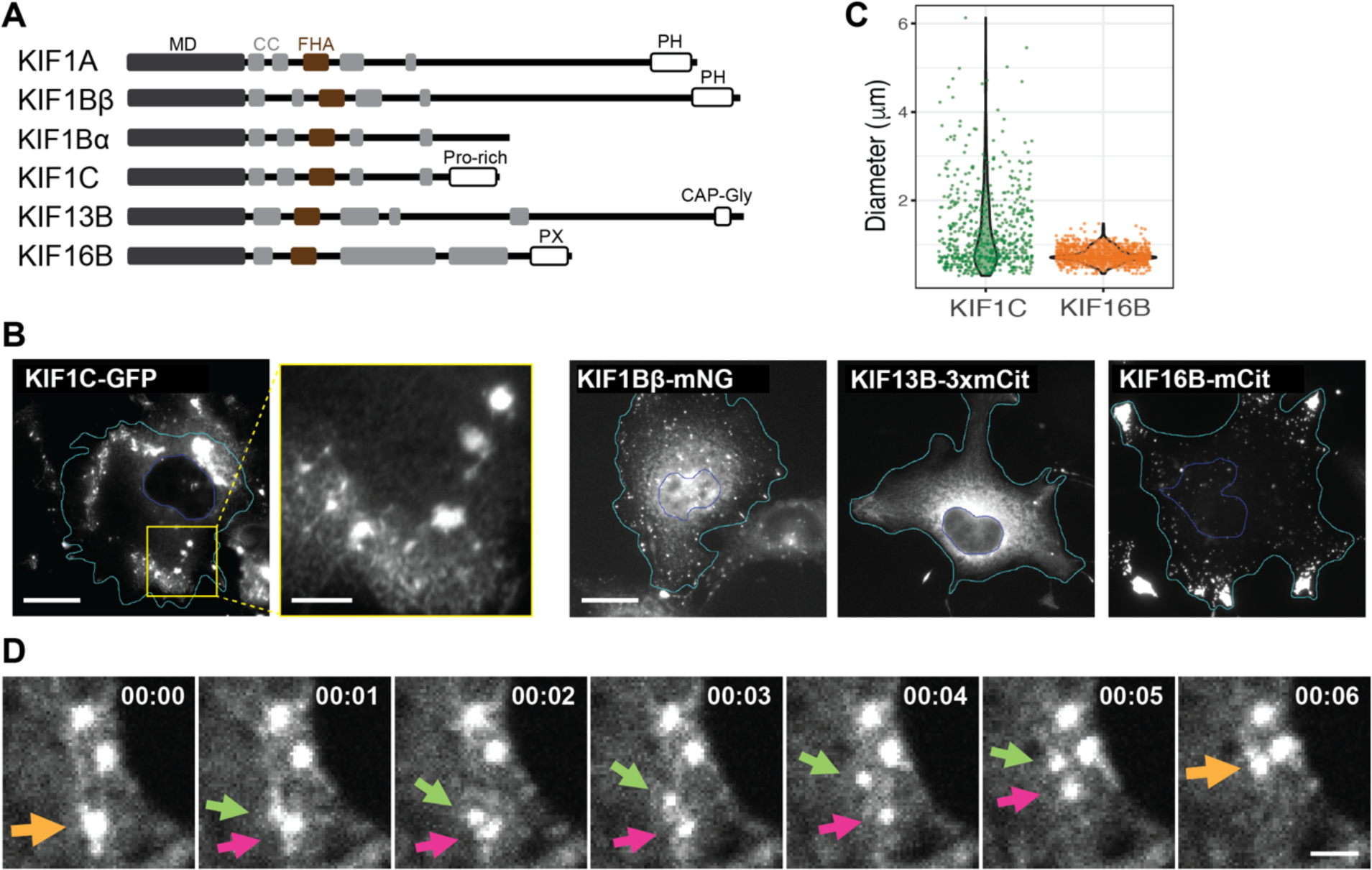
KIF1C forms dynamic puncta in cells. (A) Schematic of domain organization of the indicated kinesin-3 family members. MD: <u>M</u>otor <u>d</u>omain; CC: <u>C</u>oiled- <u>c</u>oil domain; FHA: <u>F</u>ork<u>h</u>ead-<u>a</u>ssociated domain. PH: <u>P</u>leckstrin <u>h</u>omology domain; Pro-rich: Proline-rich domain; CAP-Gly: <u>Gly</u>cine-rich domain of <u>C</u>ytoskeleton-<u>a</u>ssociated <u>p</u>roteins; PX: <u>P</u>ho<u>X</u> homology domain. (B) Localization of fluorescently-tagged kinesin-3 motors in COS-7 cells. Scale bar: 20 μm for whole cell views, 5 μm for magnified image of the KIF1C-expressing cell. Cyan lines indicate cell boundaries. Blue lines indicate nuclear boundaries. (C) Quantification of size distribution of individual KIF1C or KIF16B puncta. Each dot represents one punctum. N = 38 cells for KIF1C (601 puncta); N = 34 cells for KIF16B (1120 puncta). (D) Representative live-cell imaging showing dynamic fusion and fission events of KIF1C puncta. Scale bar: 2 μm. Time label is [min:sec].

To probe the identity of the KIF1C puncta, we stained cells for various markers but found that the KIF1C puncta did not colocalize with markers of endosomes, lysosomes, focal adhesions, or mitochondria (Fig. S1). We next examined the protein sequence of KIF1C. Like other kinesins involved in cargo transport, full-length (FL) KIF1C contains an N-terminal motor domain (MD), a stalk domain with several coiled-coil (CC) segments, and a unique C-terminal tail domain for binding specific cargoes (Fig. 1 A). KIF1C’s tail domain was previously referred to as a proline- rich domain^56,57^ and analysis of the amino acid composition of the KIF1C IDR (amino acids 886- 1103) reveals low sequence complexity, with significant enrichment not only of proline (20.2%) residues, but also arginine (13.3%), glycine (9.2%), and serine (8.7%) residues (Fig. S2 A,B).

Structure prediction using IUPred suggests that the tail domain is an intrinsically disordered region (IDR) and PLAAC analysis indicates that a subregion of the IDR is a prion-like domain (PLD) (Fig. 2 A), a type of low complexity region frequently found in RNA-binding proteins (RBPs)^58,59^. Notably, with the exception of related kinesin-3 KIF1Ba (see Discussion), long stretches of IDR with PLD are not found in the tail domains in other kinesin-3 motors or of kinesin-1 motors (Fig. S2 C,D).

**Figure 2.**
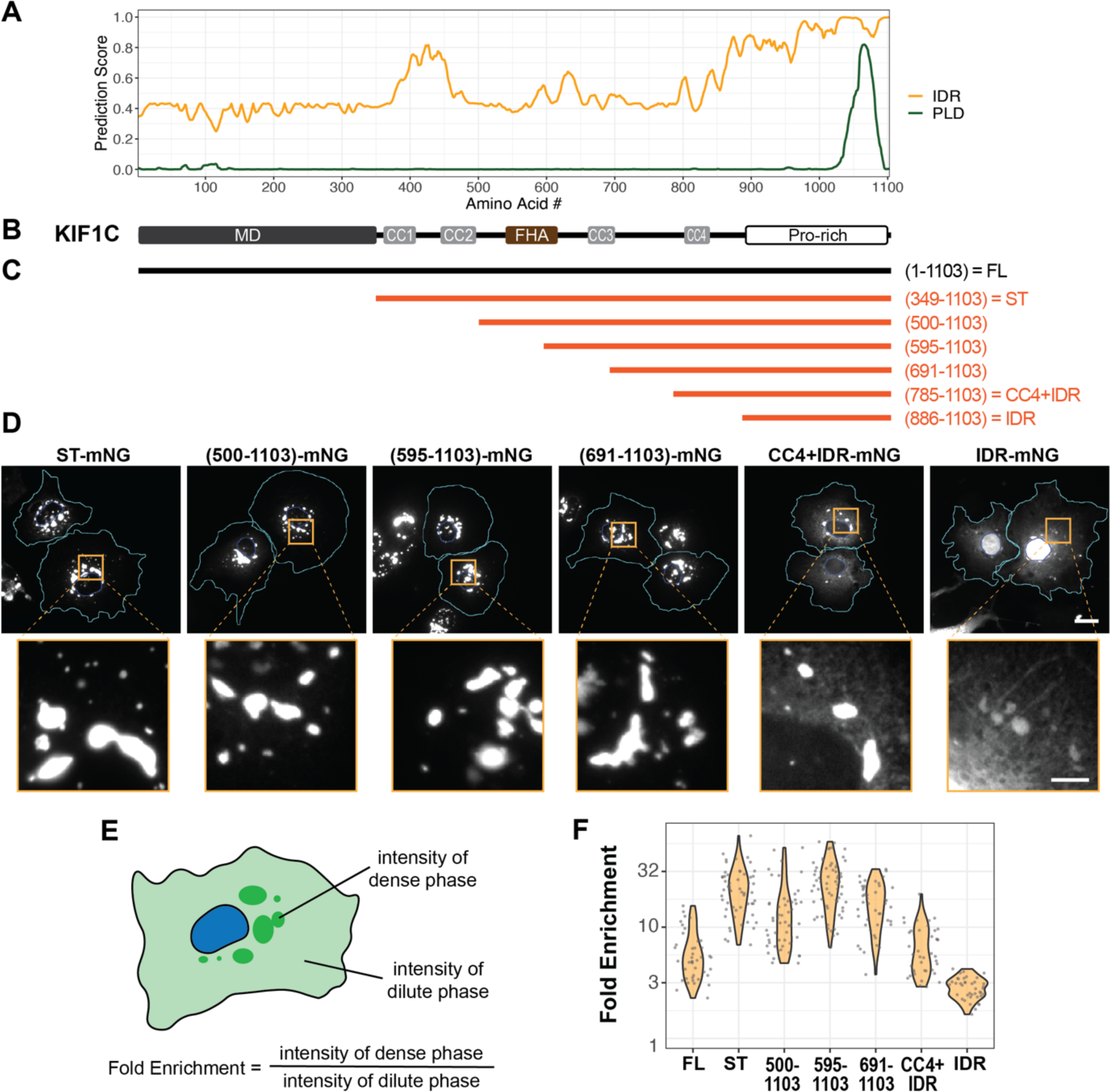
**The C-terminal IDR domain determines KIF1C puncta formation.** (A) IUPred prediction of intrinsically disordered region (IDR) and PLAAC prediction of prion-like domain (PLD) for KIF1C. x-axis: KIF1C amino acid residue number; y-axis: predicted probability of the given residue being part of an IDR (orange line) or a PLD (green line). (B) Schematic of KIF1C domain organization, aligned with the x-axis of (A). (C) Schematic of KIF1C serial truncations from the N-terminus, aligned with (B). (D) Localization of KIF1C truncations in COS-7 cells. Scale bar: 20 μm for whole cell views, 5 μm for magnified images. Cyan lines indicate cell boundaries. Blue lines indicate nuclear boundaries. (E) Schematic for quantification of the fold enrichment of KIF1C in puncta. (F) Fold enrichment of full-length or truncated KIF1C. y-axis: fold enrichment plotted on log scale. Fold enrichments: 6.0 ± 3.2 (mean ± SD) for FL, N = 42 cells; 22.7 ± 11.1 for ST, N = 58 cells; 14.2 ± 10.5 for (500-1103), N = 54 cells; 26.0 ± 12.7 for (595-1103), N =65 cells; 16.7 ± 7.5 for (691-1103), N = 45 cells; 6.8 ± 3.6 for CC4+IDR, N = 40 cells; 2.9 ± 0.6 for IDR, N = 40 cells.

To test whether the IDR is required for the formation of the KIF1C puncta, we generated KIF1C variants with serial truncations from the N-terminus (Fig. 2 C) or the C-terminus (Fig. S3 A). All constructs were tagged by mNeonGreen (mNG) and their localization was examined upon transient expression in cultured cells. Deletion of the IDR from the C-terminus of KIF1C(ΔIDR) resulted in a largely diffuse localization and further truncations also failed to form puncta in cells (Fig. S3 B). Although the ΔIDR construct did show small accumulations at the cell periphery, these appeared to be linear-shaped accumulations along microtubules, similar to a constitutively-active KIF1C construct containing only the motor domain (amino acids 1-348) (Fig. S3 C).

Deletion of the N-terminal motor domain resulted in a stalk+tail (ST) construct (amino acids 349-1103) that forms puncta localized in the perinuclear region (Fig. 2 D). Thus, the motor domain is not required for the formation of KIF1C puncta but plays a role in their cytoplasmic distribution. Deletion of increasing lengths of the stalk domain had little to no effect on the formation of the puncta or on their morphology or distribution (Fig. 2 D). Indeed, the IDR in the tail domain is sufficient for puncta formation as a construct containing only the IDR (amino acids 886-1103) formed puncta in the cytoplasm (Fig. 2 D). The IDR construct could also enter the nucleus and form puncta inside the nucleus, although the puncta in nuclei were smaller and harder to observe due to higher background intensity of diffusive IDR (Fig. S7 B). Quantification confirmed that the IDR is sufficient for punctum formation as the KIF1C(IDR) protein displayed an enrichment of mNG fluorescence intensity in the puncta over the cytoplasm (Fig. 2 E,F).

However, the coiled-coil segments in the stalk domain appear to facilitate puncta formation as the addition of increasing amounts of coiled coil resulted in increased KIF1C enrichment in puncta as compared to the IDR alone (Fig. 2 F). Taken together, these results indicate that the C-terminal IDR is necessary and sufficient for KIF1C puncta formation in the cytoplasm.

### KIF1C puncta show properties of biomolecular condensate in cells

The fact that formation of KIF1C puncta requires an IDR and is tuned by structured regions suggests that the puncta may be biomolecular condensates formed through the process of liquid-liquid phase separation (LLPS)^25,60,61^. Previous work has shown that phase separation is typically mediated by IDR or low-complexity domains that provide weak, multivalent interactions to allow molecules to stick together against entropy. Other features of the KIF1C puncta, such as their heterogenous size and their active fusion and fission, support the idea that they may be membrane-less, liquid condensates. We thus tested the hypothesis that KIF1C undergoes IDR-mediated LLPS by probing the physical and chemical properties of KIF1C puncta in cells.

We first used <u>f</u>luorescence <u>r</u>ecovery <u>a</u>fter <u>p</u>hotobleaching (FRAP) to test whether there is active exchange of molecules between KIF1C puncta (the dense phase) and the cytoplasm (the dilute phase). mCherry(mCh)-tagged full-length (FL), stalk+tail (ST), and IDR constructs (Fig. 3 A) were expressed in cells together with freely diffusive GFP as a control. Individual puncta were selected and photobleached at high laser power and the fluorescence recovery in the bleached area was measured. All three KIF1C constructs showed rapid fluorescence recovery with average half-time (τ_1/2_) of 3.5 sec for KIF1C(FL), 17.0 sec for KIF1C(ST), and ≤ 1.0 sec for KIF1C(IDR) (Fig. 3 B,C, Video 3, Video 4, Video 5). The behavior of KIF1C(IDR) was similar to that of GFP (τ_1/2_ ≤ 0.8 sec), although for both proteins, the measured τ_1/2_ values are likely an overestimation as fluorescence recovery occurred before post-bleaching images could be acquired. These results indicate that KIF1C molecules can rapidly exchange between the inside and outside of the condensate.

**Figure 3.**
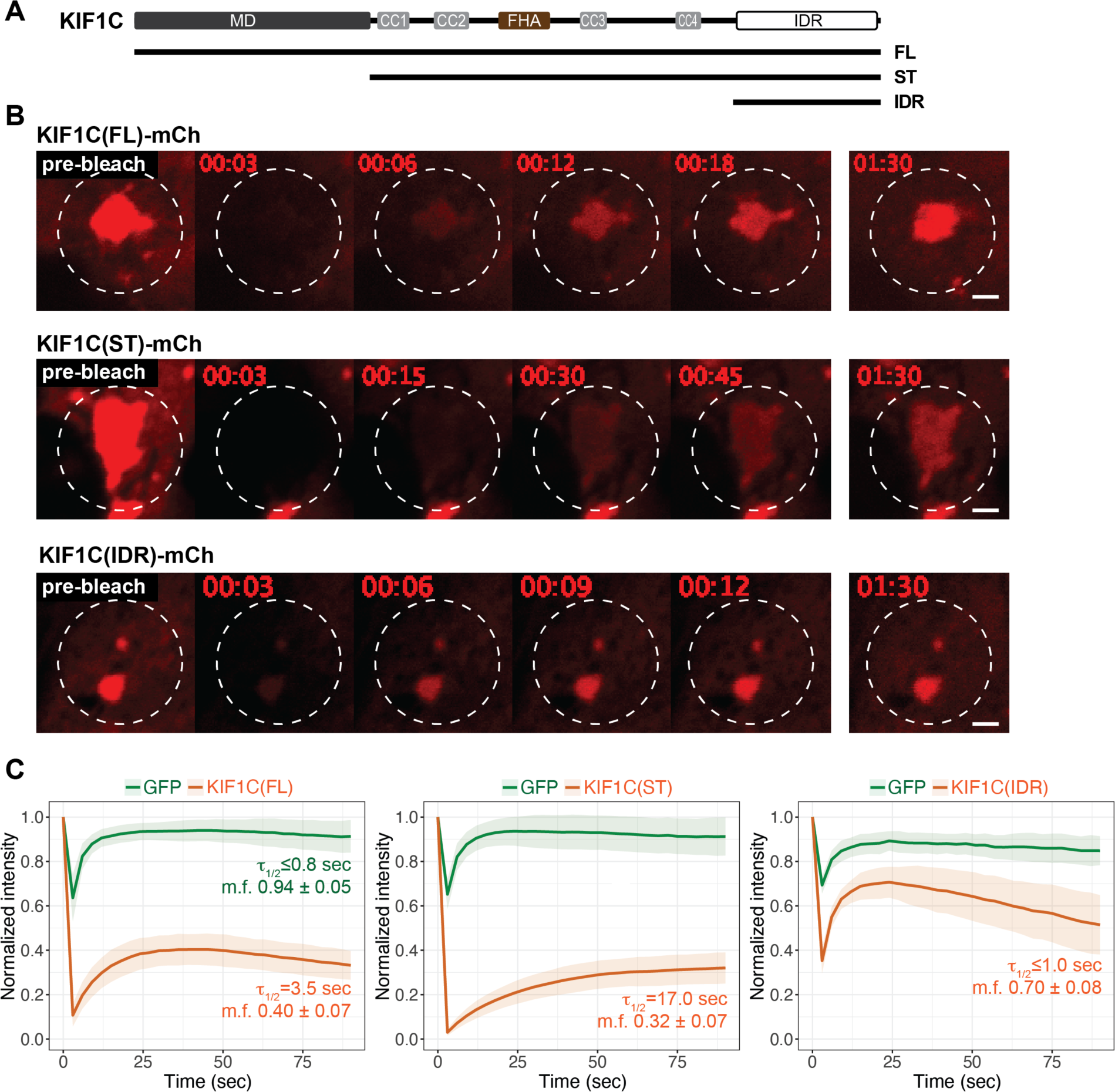
**KIF1C molecules exchange rapidly between puncta and the cytoplasm.** (A) Schematic of constructs KIF1C(FL), KIF1C(ST), and KIF1C(IDR). (B) Representative images from live-cell imaging of mCherry-tagged KIF1C constructs expressed in COS-7 cells before and after photobleaching. White dashed lines indicate photobleached regions. Scale bar: 2 μm. Time label is [min:sec]. (C) Quantification of fluorescence intensity before and after photobleaching, normalized against the frame right before photobleaching (mean ± STD). For each cell, the fluorescence recovery of co-expressed EGFP was measured as a control. N = 28 cells for KIF1C(FL); N = 20 cells for KIF1C(ST); and N = 9 cells for KIF1C(IDR). The quantified recovery half-time (τ_1/2_) and mobile fraction (m.f.) are indicated in each plot.

We next tested whether the appearance of the KIF1C puncta responds to a change in molecular concentration. To do this, we utilized a hypotonic shock assay to rapidly manipulate the cytoplasmic conditions in a reversible manner^62,63^. In this assay, treatment with hypotonic media induces an increase in cytoplasmic volume and concomitant decrease in cytoplasmic protein concentrations which can be reversed upon switching the media back to isotonic conditions (Fig. 4 A). We utilized the stalk+tail construct KIF1C(ST) to prevent motor-driven mixing of KIF1C condensates. Upon application of hypotonic media, the mNG-tagged KIF1C(ST) puncta dissolved within minutes (Fig. 4 B). By live-cell imaging, two stages could be observed: an initial stage of larger puncta disassembling into smaller puncta, and a second stage of small puncta disappearing into the diffusive phase (Video 6). Strikingly, when the solution was returned to isotonic conditions, numerous small puncta appeared in the cytoplasm, and then fused into larger puncta that localized in the perinuclear region (Fig. 4 B,C). As a control, we tested whether KIF16B puncta undergo reversible dissolution and reformation under these conditions as KIF16B is known to bind directly to endosomal membranes via the PX domain in its tail domain^55^. During the iso-hypo-isotonic cycle, KIF16B puncta did not dissolve or undergo fission/fusion (Fig. S 4, Video 7). Overall, we conclude that KIF1C puncta are liquid condensates formed through LLPS.

**Figure 4.**
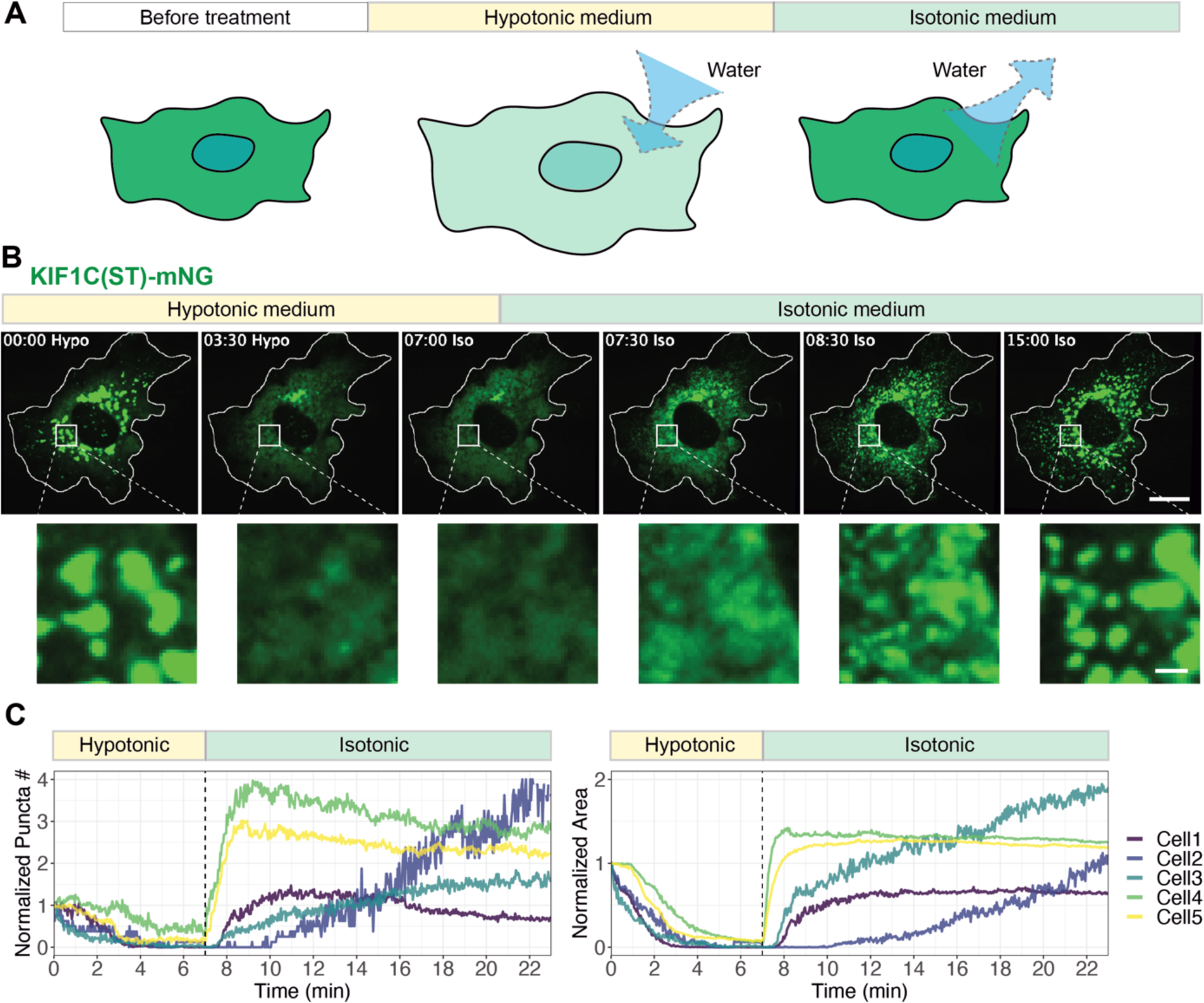
**KIF1C condensates display LLPS properties in cells.** (A) Schematic of the cytoplasm dilution assay for testing LLPS in cells. (B) Representative images of mNG-tagged KIF1C(ST) localization before treatment, during hypotonic treatment, and upon return to isotonic media. Scale bar: 20 μm for whole cell views, 2 μm for magnified images. Time label is [min:sec]. (C) Quantification of change of puncta number (left) and total area of puncta (right) over time. x axis: time, with a vertical dashed line indicating the time point of switching from hypotonic media to isotonic media. y axis: puncta number or puncta area normalized against the initial state in the first frame of live-cell imaging. The example cell in (B) is Cell 5 in the plots.

Recent work has shown that several microtubule-associated proteins can undergo LLPS and “wetting” of the microtubule surface^64–69^. CLIP-170 is of particular interest as it also undergoes LLPS at microtubule plus ends in the cell periphery^67,68^ . We thus tested whether the KIF1C condensates are related to those of CLIP-170, however no colocalization was observed between mCh-tagged KIF1C condensates and GFP-tagged CLIP-170 condensates, suggesting that their LLPS is mutually exclusive (Fig. S5 A). Several of these microtubule-associated proteins can enrich free tubulin in the condensate via their microtubule-binding domains^70–74^. We thus tested whether KIF1C condensates can enrich tubulin, however, we found that KIF1C condensates do not recruit free tubulin upon depolymerization of cytoplasmic microtubules (Fig. S5 B).

Therefore, KIF1C condensates appear to be distinct from other cytoskeletal condensates.

### The KIF1C IDR is sufficient for driving LLPS in vitro

To directly test whether KIF1C can undergo LLPS on its own or whether it requires additional components in the cytoplasm, we purified mNG-tagged KIF1C IDR (aa 886-1103) and CC4+IDR (aa 785-1103) constructs from *E. coli* (Fig. 5 A; Fig. S6 A). Both proteins were purified under high salt conditions (500 mM NaCl) and then underwent phase separation *in vitro* upon lowering the NaCl concentration to 100 mM (Fig. 5 B). Both proteins formed small droplets whereas purified mNG did not form visible structures under the same conditions (Fig. 5 B). At 2 uM protein concentration and 100 mM NaCl, the KIF1C(IDR) droplets were smaller [diameter 0.43 ± 0.22 μm (mean ± STD)] than KIF1C(CC4+IDR) droplets [0.48 ± 0.27 μm (mean ± STD)] (Fig. 5 C). The KIF1C(IDR) puncta were also less round than those of KIF1C(CC4+IDR) (Fig. 5 C). Finally, the KIF1C(CC4+IDR) condensates were observed to undergo typical fusion behavior (Fig. 5 D; Video 8, Video 9) whereas the KIF1C(IDR) puncta appeared to stick together rather than undergo fusion, even after 2 hours of incubation (Fig. S5 B). These results suggest that both IDR and CC4+IDR are sufficient to drive LLPS by themselves *in vitro*, but the CC4+IDR condensates are more liquid-like whereas the IDR condensates are more hydrogel-like.

**Figure 5.**
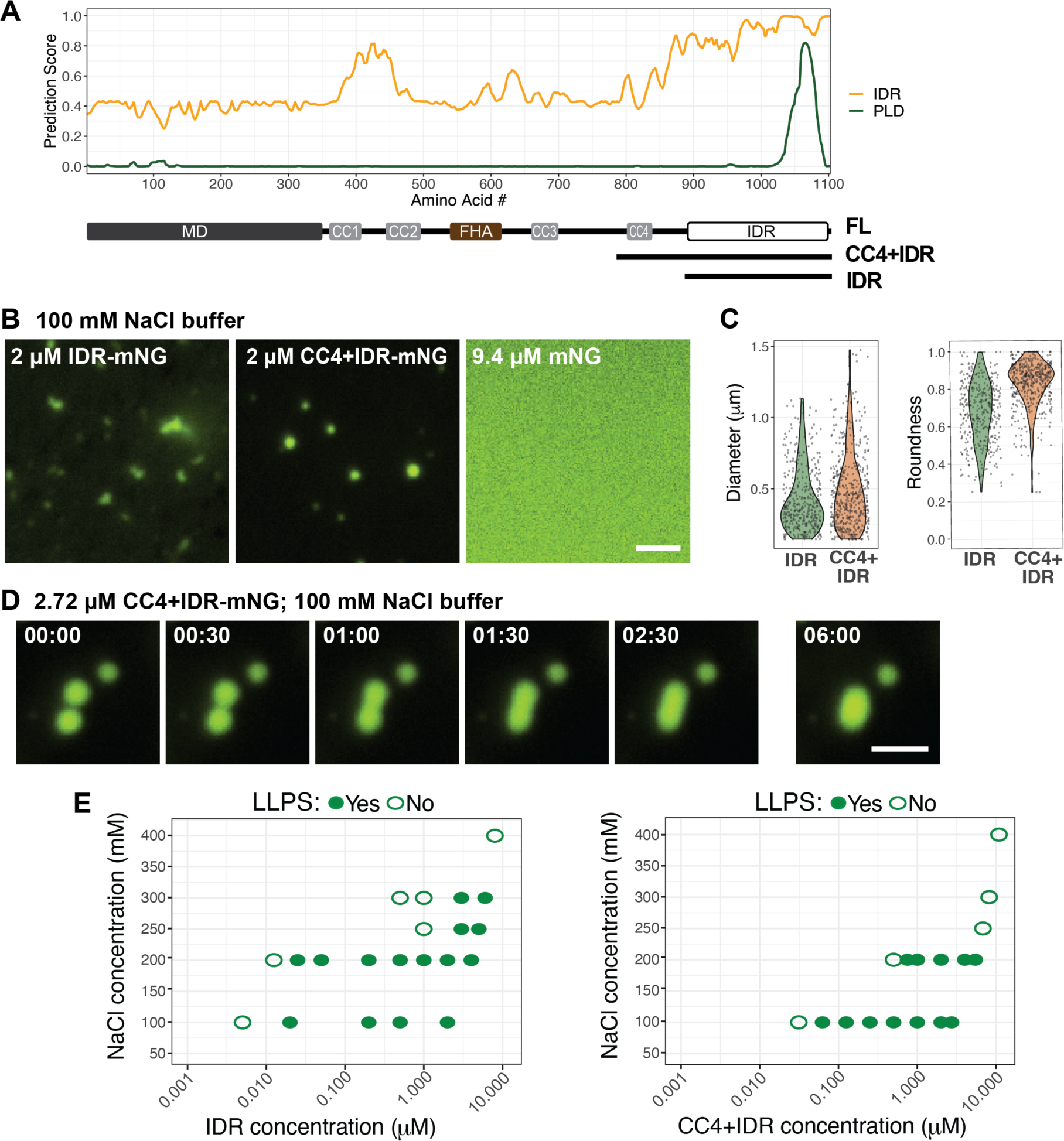
**The KIF1C IDR is sufficient for driving LLPS in vitro.** (A) Schematic of the mNG-tagged KIF1C truncations CC4+IDR and IDR. (B) Purified IDR-mNG and CC4+IDR-mNG undergo LLPS in 100 mM NaCl buffer whereas purified mNG does not. Scale bar: 2 μm. (C) Quantification of condensate size (diameter) and shape (roundness) under the conditions in (B). N=389 puncta for IDR-mNG; N=503 puncta for CC4+IDR-mNG. (D) Representative images of a fusion event observed for CC4+IDR condensates. Scale bar: 2 μm. Time label is [min:sec]. (E) Phase diagrams of condensate formation for IDR-mNG and CC4+IDR-mNG at different salt concentrations. x- axis: protein concentration plotted on log scale; y-axis: NaCl concentrations of buffer. Solid green dots indicate that LLPS **was** observed under given conditions, open green circles indicate that LLPS was **not** observed under given conditions.

We next determined the phase separation diagrams for LLPS of the CC4+IDR and IDR constructs at different NaCl concentrations. The KIF1C(IDR) was able to form condensates at higher salt concentrations (250-300 mM) than the KIF1C(CC4+IDR) construct (Fig. 5 E). In 100 mM NaCl buffer, which has an ionic strength close to the physiological range, the critical concentrations for LLPS were ∼20 nM for IDR and ∼62.5 nM for CC4+IDR (Fig. 5 E).

Interestingly, these values are close to the estimated concentration of endogenous KIF1C [25.5 ± 7.6 nM, (mean ± STD), Fig. S5 C]. Overall, these results demonstrate that the KIF1C IDR is sufficient to drive LLPS.

### KIF1C condensates enrich RNA and facilitate mRNA transport in cells

To our knowledge, KIF1C is the first kinesin reported to undergo LLPS. As a kinesin, what does LLPS activity mean for KIF1C’s function? Recently, several groups reported that KIF1C associates with and transports mRNAs^43–46^. These findings are particularly interesting in light of KIF1C’s ability to undergo LLPS, as many phase-separating proteins are known to be RNA-binding proteins (RBPs)^27,75^. Furthermore, KIF1C contains a putative prion-like domain (PLD, Fig. 2 A), a domain that is known to mediate LLPS of RBPs such as FUS and TDP-43^76–82^.

Therefore, we explored whether KIF1C condensates are related to other RNA granules and whether they contain RNAs.

To test whether KIF1C can interact with other RNA granules in cells, we overexpressed KIF1C in COS-7 cells and found that KIF1C condensates colocalize with protein markers of two types of RNA granules, stress granules and P-bodies, by immunofluorescence staining (Fig. S7 A).

The IDR is necessary and sufficient for interaction with RNA granules as the KIF1C(IDR) construct colocalized with both stress granules and P-bodies, whereas the KIF1C(ΔIDR) construct did not colocalize with either membrane-less organelle (Fig S7 A).

To directly test whether KIF1C condensates can incorporate RNA, fluorescently-labeled synthesized RNA oligos were introduced into cells by transient transfection. GU-rich RNA oligos were incorporated into KIF1C(ST) and KIF1C(IDR) condensates in cells whereas polyA RNA oligos were not incorporated into condensates formed by either construct (Fig. 6 A), suggesting that there is sequence selectivity for RNA incorporation into KIF1C condensates. For KIF1C(IDR), incorporation of GU-rich oligos was observed in condensates in both the cytoplasm and the nucleus (Fig. S7 B).

**Figure 6.**
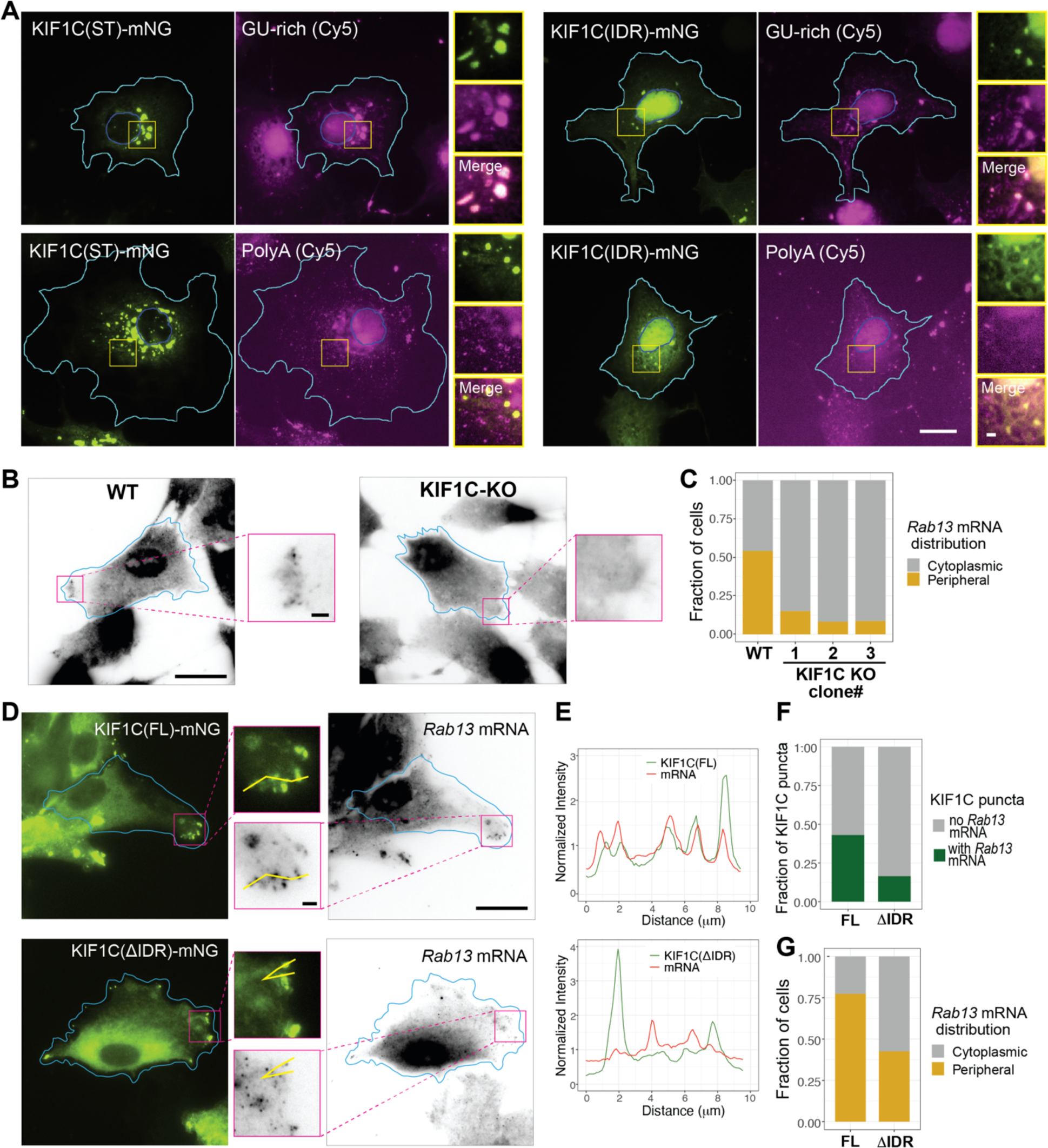
**KIF1C LLPS selectively enriches exogenous and endogenous RNA.** (A) Fluorescently-labelled (Cy5) GU-rich or polyA RNA oligos were introduced into COS-7 cells expressing KIF1C(ST)-mNG (left panels) or KIF1C(IDR)-mNG (right panels). Representative images are shown. Cyan lines indicate cell boundaries. Blue lines indicate nuclear boundaries. The yellow box indicates the region in the magnified images to the right. Scale bar: 20 μm for whole cell views, 2 μm for magnified images. (B,C) Distribution of endogenous *Rab13* mRNA revealed by smFISH in WT (left) and KIF1C-KO (right) hTERT- RPE1 cells. (B) Representative smFISH images. Scale bar: 20 μm for whole cell views, 2 μm for magnified images. Cyan lines indicate cell boundaries. (C) Quantification of the fraction of cells displaying cytoplasmic or peripheral distribution of *Rab13* mRNA in WT hTERT-RPE1 cells and three KIF1C-KO clones. N = 72 cells for WT, N = 66 cells KIF1C-KO-clone1, N = 73 cells for KIF1C-KO-clone2, and N = 69 cells for KIF1C-KO-clone3. y-axis: fraction of cells showing peripheral or cytoplasmic distribution of *Rab13* mRNA. (D-G) Rescue of *Rab13* mRNA enrichment and localization in KIF1CKO-clone1 cells. (D) Representative images of expressed KIF1C(FL)-mNG (top) or KIF1C(ΔIDR)-mNG (bottom) and smFISH for endogenous *Rab13* mRNA. Cyan lines indicate cell boundaries, magenta box indicates the region of the magnified imaged in the middle. Yellow lines in the magnified images indicate line scan for intensity profiles in (E). Scale bar: 20 μm for whole cell views, 2 μm for magnified images. (E) Fluorescence intensity profiles of KIF1C protein (green lines) and *Rab13* mRNA (red lines). x-axis, distance along the scanning line. y-axis, intensity values normalized to the average intensity along the entire line. (F) Quantification of the fraction of cells with peripheral or cytoplasmic distribution of *Rab13* mRNA in cells expressing KIF1C(FL)-mNG (N = 84 cells) or KIF1C(ΔIDR)-mNG (N = 68 cells). (G) Quantification of the fraction of KIF1C puncta with or without *Rab13* mRNA enrichment for KIF1C(FL)-mNG (N = 64 cells) and KIF1C(ΔIDR)-mNG (N = 63 cells).

To test whether the KIF1C IDR and LLPS are required for the localization of endogenous mRNAs, we generated KIF1C knockout (KO) hTERT-hRPE1 cells using CRISPR-Cas9 gene editing. Three cell lines were selected and all three contained a frameshift mutation in amino acid 6, resulting in a premature stop codon (Fig. S8 A) and undetectable protein levels by western blotting (Fig. S8 B). We confirmed that all three KIF1C KO clones display a defect in the distribution of *Rab13* mRNA to cell protrusions, as visualized by <u>s</u>ingle-<u>m</u>olecule <u>f</u>luorescence *in <u>s</u>itu* <u>h</u>ybridization (smFISH) (Fig. 6 B,C), consistent with previous studies using RNAi to deplete KIF1C expression^43^.

We then expressed KIF1C(FL) and KIF1C (IDR) in KIF1C KO cells. KIF1C(FL) localized diffusely in cells and in puncta at the cell periphery (Fig. 6 D), due to its ability to undergo microtubule-based motility and LLPS. The KIF1C(FL) condensates showed strong colocalization with *Rab13* mRNA (Fig. 6 E,F) and were able to efficiently rescue *Rab13* mRNA localization to the cell periphery (Fig. 6 G) In contrast, although KIF1C(ΔIDR) localized to the cell periphery due to its ability to undergo microtubule-based motility, it showed less colocalization with *Rab13* mRNA (Fig. 6 E,F) and was less efficient at rescuing *Rab13* mRNA at the cell periphery (Fig. 6 G). Together, these data indicate that KIF1C condensates can incorporate exogenous and endogenous RNAs and that KIF1C is involved in the targeting of select mRNAs to the cell periphery.

### A subregion of the KIF1C IDR critical for enrichment of Rab13 mRNA in condensates

These results also indicate that the IDR is responsible for both LLPS and incorporation of select RNAs. To test whether KIF1C LLPS and RNA binding are interdependent or two separate processes, we sought to distinguish sequences for LLPS and RNA binding within the IDR. We first hypothesized that the prion-like domain (PLD), a type of low complexity region frequently found in RNA-binding proteins (RBPs)^58,59^ would be involved in enrichment of *Rab13* mRNA into the condensates. However, deletion of the PLD [ST(DPLD), deletion of amino acids 1054- 1080] or mutation of all glutamine and asparagine amino acids to alanines [ST(PLDmut)] (Fig. 7 A) had no effect on condensate formation or the incorporation of *Rab13* mRNA (Fig. 7 B, E).

**Figure 7.**
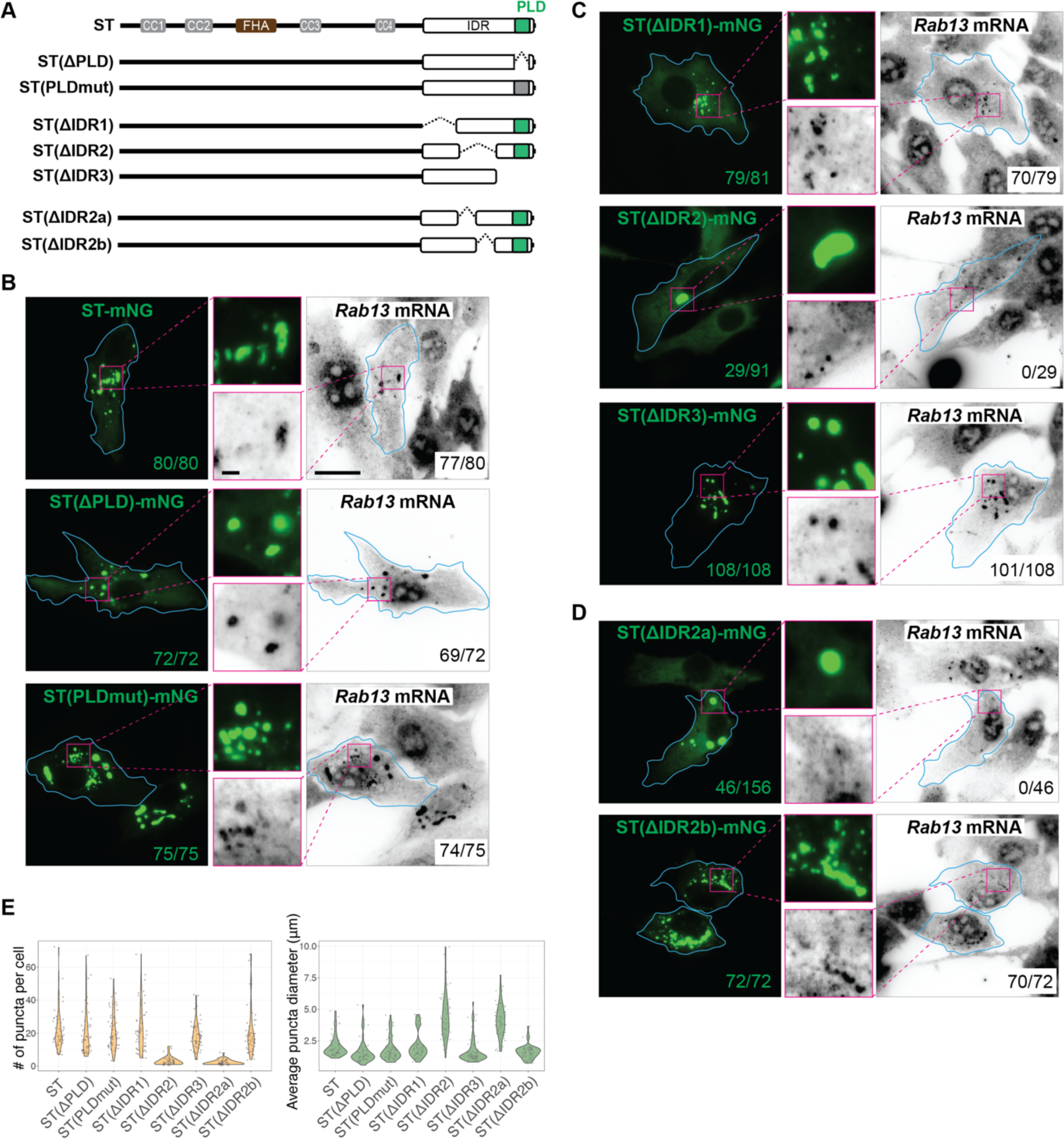
**KIF1C IDR subregion required for Rab13 mRNA localization.** (A) Schematic of constructs KIF1C(ST), ST(ΔPLD), ST(PLDmut), ST(ΔIDR1), ST(ΔIDR2), ST(ΔIDR3), ST(ΔIDR2a), and ST(ΔIDR2b). The PLDmut construct has 1059N, 1063Q, 1066Q, and 1071Q all mutated to A. (B-D) Localization of *Rab13* mRNA in KIF1CKO-clone 1 cells expressing (B) deletion or mutation of the PLD, (C) deletion of subregions of the IDR, or (D) deletion of subregions IDR2a or IDR2b. Cyan lines indicate cell boundaries; magenta boxes indicate the region in the magnified images in the middles. Scale bar: 20 μm for whole cell views, 2 μm for magnified images. Numbers in green text: number of cells with puncta; numbers in black text: number of cells with *Rab13* mRNA enrichment in the KIF1C condensates. (E) Quantification of puncta number (left) and puncta diameter (right) for each construct. Each dot represents one cell. For ST: 21.8 ± 12.0 puncta per cell (mean ± STD), 2.06 ± 0.83 μm diameter (mean ± STD), N = 49 cells; For ST(ΔPLD): 20.1 ± 13.7 puncta per cell, 1.62 ± 0.94 μm diameter, N= 47 cells; For ST(PLDmut): 20.8 ± 11.1 puncta per cell, 1.83 ± 0.89 μm diameter, N = 56 cells; For ST(ΔIDR1), 23.1 ± 14.3 puncta per cell, 2.13 ± 1.04 μm diameter, N = 53 cells; For ST(ΔIDR2), is 3.1 ± 2.4 puncta per cell, 4.44 ± 1.74 μm diameter, N = 51 cells; For ST(ΔIDR3), 18.4 ± 8.1 puncta per cell, 1.71 ± 0.98 μm diameter, N = 62 cells; For ST(ΔIDR2a), 2.4 ± 1.5 puncta per cell, 4.15 ± 1.28 μm diameter, N = 51 cells; For ST(ΔIDR2b), 20.9 ± 14.1 puncta per cell, 1.63 ± 0.55 μm diameter, N = 53 cells.

We thus generated larger deletions across the IDR region (Fig. 7 A). Deleting the first or last third of the IDR [constructs ST(DIDR1) and ST(DIDR3), respectively; Fig. 7 A] had no effect on condensate formation or *Rab13* mRNA incorporation (Fig. 7 C,E). In contrast, deleting the middle third of the IDR [construct ST(DIDR2), Fig. 7 A] altered the ability to undergo LLPS, with condensates forming in only ∼30% of the cells and the condensates that did form appearing fewer in number and larger in size (Fig. 7 C,E). Importantly, the ST(DIDR2) condensates that did form were unable to incorporate *Rab13* mRNA (Fig. 7 C). This result suggests that the IDR-driven LLPS of KIF1C does not depend on mRNA incorporation, but is strongly affected by it. These results also show that the LLPS is resistant to truncations of large portions of IDR, consistent with the theory that LLPS is driven by many multivalent interactions along the IDR, rather than specific interaction sites.

To further narrow down the RNA-binding region of the KIF1C IDR, we generated two further deletions of IDR2. Deleting the first half of IDR2 [ST(DIDR2a), Fig. 7 A] resulted in the formation of fewer and larger condensates in the expressing cells and a failure to incorporate *Rab13* mRNA (Fig. 7 D,E), similar to the full IDR2 deletion. In contrast, deleting the second half [ST(DIDR2b), Fig. 7 A] did not affect condensate formation or mRNA incorporation (Fig. 7 D,E). These results identify a segment of 47 amino acids (IDR2a, amino acids 938-985) to be the region responsible for RNA incorporation into KIF1C condensates.

### The LLPS behavior of KIF1C is buffered by the cytoplasmic RNA pool

Previous work has suggested that LLPS of prion-like RBPs, such as FUS, TDP-43, and TAF15, is buffered by the non-specific RNA pool in the cytoplasm and/or nucleoplasm^83,84^. Given KIF1C’s PLD, we tested whether its LLPS is also sensitive to the non-specific RNA pool, we microinjected RNase A to remove the cytoplasmic RNA in a non-specific manner. We again used the KIF1C(ST) construct to prevent motor-driven movement along microtubules and carried out live-cell imaging before and for ∼20 min after RNase A microinjection. After injection of RNase A, existing KIF1C(ST) condensates showed no change in morphology or distribution but strikingly, new KIF1C(ST) puncta appeared within minutes (Fig. 8 A; Video 10). The new condensates were dynamic and often underwent fusion to make larger puncta. In contrast, the injection marker rhodamine-dextran did not alter the KIF1C(ST) condensates (Fig. 8 B; Video 11). These results demonstrate that KIF1C’s LLPS behavior is sensitive to the endogenous cytoplasmic RNA content, and supports the idea that KIF1C’s interaction with RNA molecules is sequence selective, similar to other prion-like RBPs. Together with the effects of IDR2a deletion (Fig. 7), the RNase A injection results provide evidence that RNA incorporation modulates the LLPS behavior of KIF1C.

**Figure 8.**
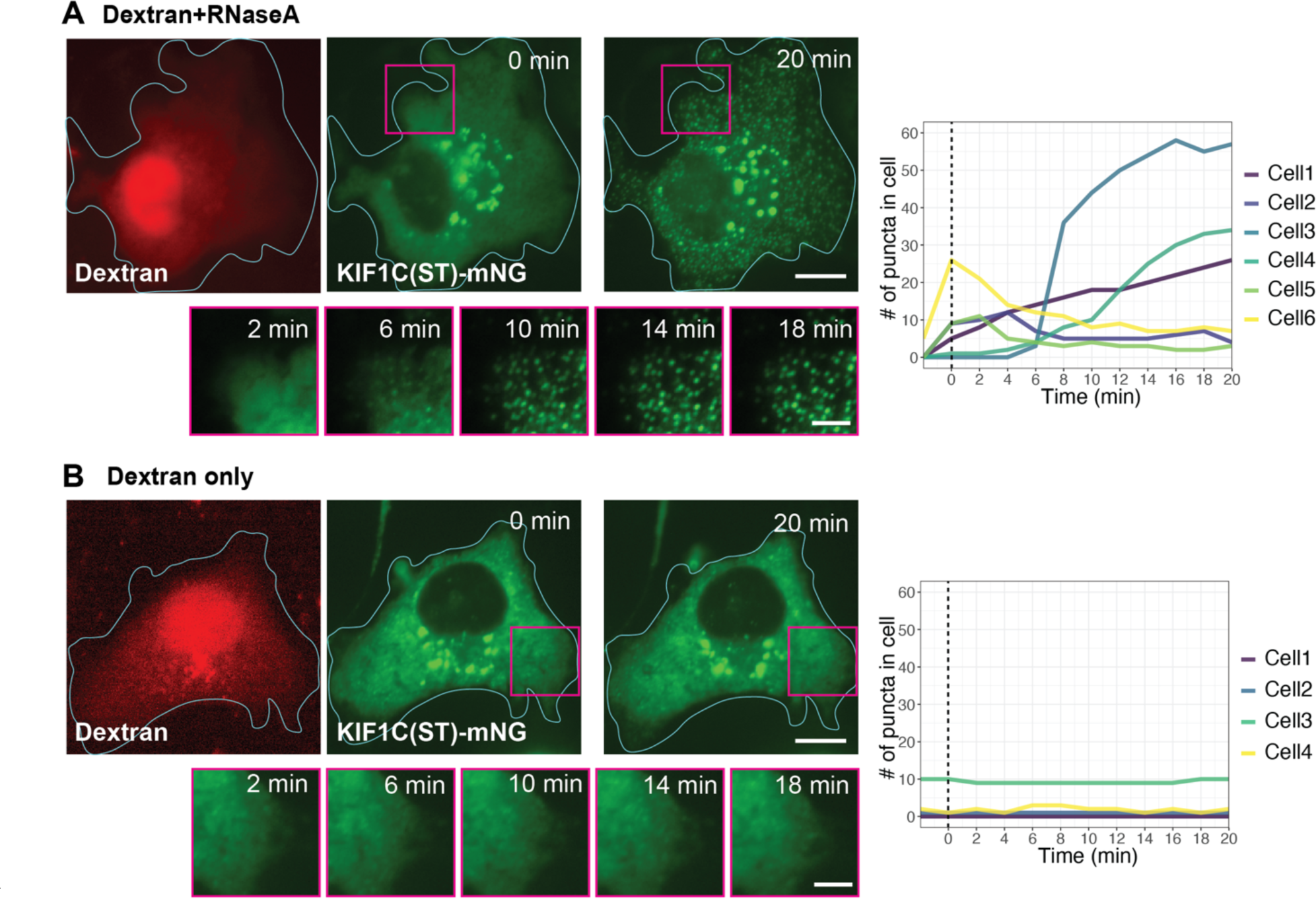
The cytoplasmic RNA pool buffers KIF1C LLPS. (A,B) Representative images of a COS-7 cell expressing KIF1C(ST)-mNG before and after microinjection with (A) RNaseA and fluorescent dextran or (B) fluorescent dextran only. Cyan lines indicate cell boundaries. Scale bar: 20 μm for whole cell views, 5 μm for magnified images. The graphs on the right indicate quantification of the number of puncta in the red-boxed areas over time. y-axis, number of KIF1C-ST puncta. x-axis, time after injection. The vertical dashed lines represent the timepoint of microinjection.

## Discussion

Our findings demonstrate that the kinesin-3 member KIF1C undergoes LLPS to form biomolecular condensates in the cytoplasm of mammalian cells. The IDR in the KIF1C tail domain is required for LLPS and the physical properties of the resulting condensate are modulated by the adjacent CC4 structural domain. KIF1C’s LLPS behavior enables it to function in RNA localization via selective recruitment of mRNA molecules into the KIF1C condensates. Given previous work on KIF1C’s roles in mRNA transport and cell migration^16,43,85^, we propose that KIF1C drives microtubule-based transport of specific mRNAs to the cell periphery and that its accumulation at microtubule plus ends enables the formation of biomolecular condensates that accumulate mRNAs and regulate the concentration and/or activity of the encoded proteins at cell protrusions (Fig. 9). Future investigations will help us understand how KIF1C LLPS regulates the binding of specific mRNAs and their processing, translation, and stability in cells.

**Figure 9.**
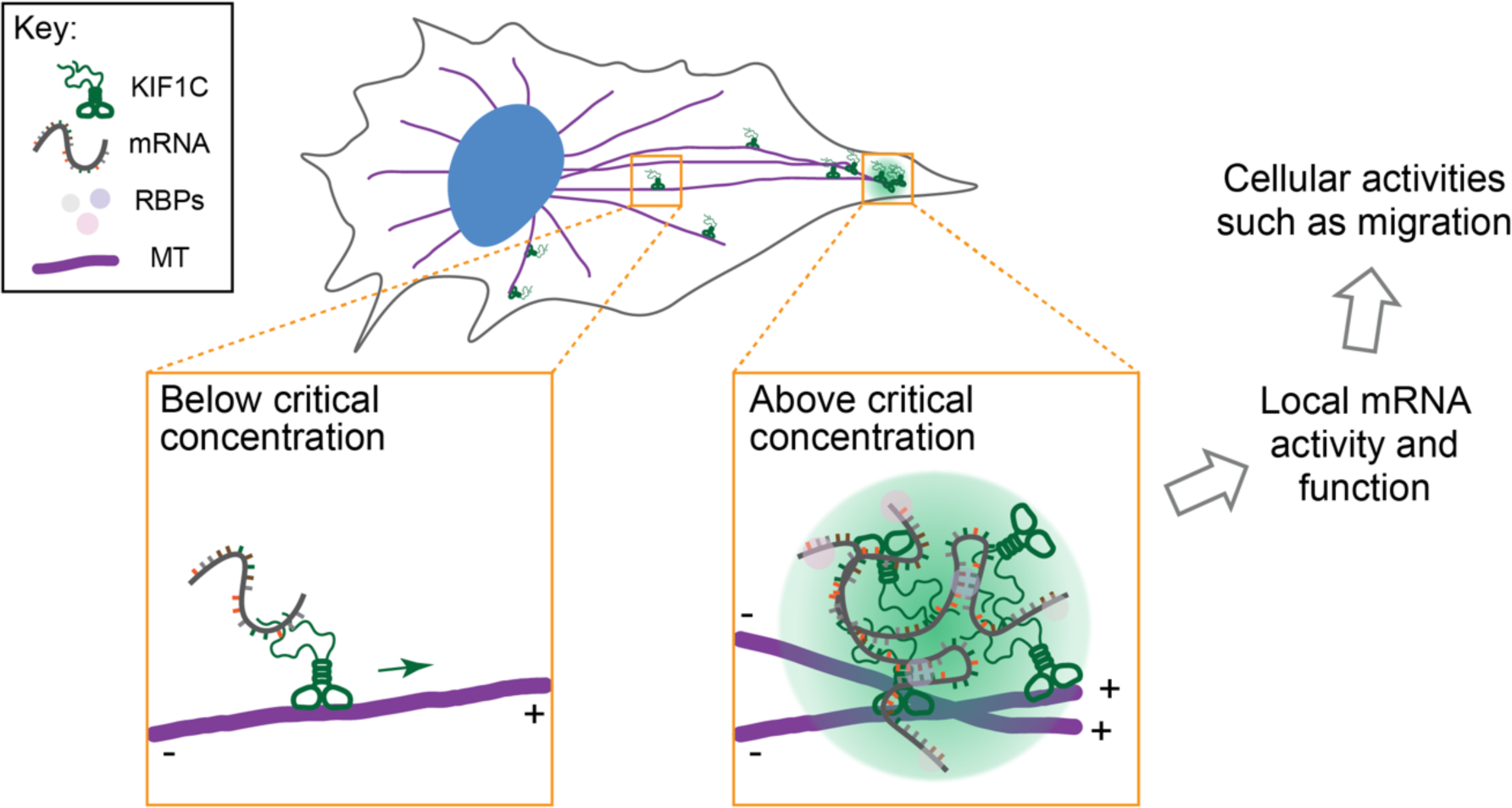
Working model of KIF1C condensate formation during mRNA transport. KIF1C drives microtubule-based transport of specific mRNAs to the cell periphery, largely as single motors. Upon reaching the microtubule plus ends in the cell periphery, KIF1C accumulates above the critical concentration of LLPS, driving the formation of biomolecular condensates that regulate mRNA activity and function during polarized cellular processes such as migration.

### LLPS of kinesins and other microtubule-associated proteins

We demonstrate that the IDR in the C-terminal tail domain of KIF1C is necessary and sufficient for LLPS and formation of a biomolecular condensate with liquid-like properties. KIF1C is the first kinesin, to our knowledge, demonstrated to undergo LLPS despite the fact that many of the 45 kinesin genes in the human genome contain significant regions of intrinsically disordered residues^86^. Within the kinesin-3 family, KIF14 contains a ∼350 amino acid segment N-terminal to its motor domain which is predicted to be intrinsically disordered. Although phase separation of the purified KIF14 IDR has not been tested, the KIF14 IDR does not form puncta when expressed in mammalian cells^87^, suggesting that it does not undergo LLPS. Rather, KIF14’s IDR has been shown to bind partner proteins such as PRC1 and actin and to regulate KIF14 motility^88,89^.

Of the other kinesin-3 family members, only the KIF1Bα tail domain shows a high score in IDR and PLD predictions (Fig S2 C). KIF1Bα is closely related to KIF1C as they share ∼62% amino acid sequence identity overall, and ∼33% identity within their IDR domains (KIF1Bα amino acids 870-1153, KIF1C amino acids 886-1103). A KIF1Bα stalk+tail construct, KIF1Bα(ST), forms small puncta when expressed in hTERT-RPE cells but forms a network-like structure when expressed in COS-7 cells (Fig S8 A), suggesting that KIF1Bα may also be capable of LLPS. However, KIF1Bα puncta do not associate with RNA-containing P-bodies and stress granules (Fig S9 B), recruit exogenous RNA oligos (Fig S9 C), or recruit endogenous *Rab13* mRNA (Fig S9 D).

Microtubules have emerged as a platform for LLPS^69^ as several non-motor microtubule-associated proteins (MAPs) can undergo LLPS, including tau, EB1, CLIP-170, BuGZ, TPX2, and CAMSAP2, and thereby regulate microtubule nucleation, branching, bundling, and/or mitotic spindle formation^64–68,70–72,90–93^. However, KIF1C condensates appear to be distinct from these other microtubule-associated biomolecular condensates. For example, although both KIF1C and CLIP-170 undergo LLPS at microtubule plus ends, KIF1C condensates do not recruit free tubulin (Fig S5), while RNA molecules are excluded from CLIP-170 condensates^67^.

### The KIF1C IDR and coiled-coil domains influence the rheological properties of the condensate

We demonstrate that the C-terminal KIF1C IDR is essential for LLPS as constructs that lack the IDR do not form droplets in mammalian cells whereas constructs that contain only the IDR form droplets in cells and *in vitro*. We find that the purified IDR protein readily undergoes LLPS at physiological salt conditions, in the absence of crowding agents, and near the estimated KIF1C endogenous concentration. This is in contrast to other MAPs that accumulate at microtubule plus ends, such as EB1 and CLIP-170, which undergo LLPS at mM concentrations or in the presence of crowding agents^68,68,92,93^.

Several aspects of KIF1C’s IDR and resulting LLPS are reminiscent of findings on prion-like RBPs, such as FUS, TDP-43, and TAF15. First, KIF1C and the prion-like RBPs show robust IDR-dependent phase separation in the absence of crowding agents at low protein concentrations and physiologically-relevant salt concentrations^94–96^. Second, the RBD of KIF1C and the prion- like RBPs^97^ is involved in both phase separation and RNA binding, suggesting that RNA can regulate phase separation. Consistent with this, the LLPS behavior of KIF1C and prion-like RBPs is buffered by the non-selective RNA pool^83,84^. Third, KIF1C and the prion-like RBPs share sequence similarity in their IDRs. For the prion-like RBPs, phase separation derives from collective interactions between R residues in the RBD and Y residues in the PLD^96^. For KIF1C, the RBD (IDR2a) is enriched in R residues (Fig. S10 A) and is required for phase separation whereas the PLD (in IDR3) is enriched in Y residues (Fig S10 A) but is dispensable for phase separation. Future studies are needed to understand the interplay between RNA, the RBD, and the PLD in regulating the phase behavior of KIF1C.

We also find that although the C-terminal KIF1C IDR contains the requisite sequence features to drive phase separation, inclusion of the adjacent structured CC4 segment can tune the rheological properties of the resulting condensates. *In vitro*, droplets formed by purified KIF1C IDR protein are irregular in shape and do not undergo fusion whereas droplets formed by purified KIF1C(CC4+IDR) protein are round in shape, undergo fission and fusion, and require a higher concentration for phase separation. These results suggest that KIF1C(CC4+IDR) condensates more liquid-like, indicating that inclusion of the structured domain makes the condensate more fluid. These findings indicate that LLPS of KIF1C fits within the theory of associative polymers, or the sticker-and-spacer theory, in which the IDR is mostly composed of the associative motifs, or stickers, that provide the multivalency for molecular interactions and is the main component determining the phase separation behavior whereas structured domains like CC4 act as spacers, which impact the flexibility of molecules and tune the rheological properties of condensates^98,99,96,100^.

### Mechanisms of mRNA binding by kinesins

RNA transport requires microtubule-dependent motor proteins and several kinesin family members in addition to KIF1C have been implicated^4,101,102^. An outstanding question in the field is the molecular mechanisms by which motors associate with mRNA and other RNAs. One possibility is that kinesins can bind directly to RNAs. For KIF1C, this possibility is supported by recent work demonstrating that KIF1C can undergo direct binding to RNAs in the EJC^44^.

Furthermore, KIF1C was the only kinesin identified in two studies that used UV crosslinking methods to capture RNA-binding proteins in an unbiased manner^51,52^.

An alternative possibility is that RBPs serve as adaptor proteins to link kinesins to specific mRNA cargoes in cells. This possibility fits with the classical kinesin-adaptor-cargo model^33^ and has been shown for kinesin-1 and kinesin-2 which utilize adapter proteins to link to mRNA cargoes^103,104^. For KIF1C, this possibility is supported by recent work demonstrating that muscle- blind like (MBNL) serves as an adaptor protein bridging KIF1C and mRNAs^46^. It is important to note that these models of direct RNA binding and adaptor-mediated RNA binding are not mutually exclusive and a hybrid model is also possible in which RBPs stabilize the direct interaction of kinesins with specific mRNAs, as has been proposed for kinesin-1 and the adaptor protein atypical tropomyosin 1^105,106^.

### Relationship between KIF1C LLPS and mRNA transport

We show that the KIF1C IDR is required for both LLPS and delivery of *Rab13* mRNA to the cell periphery. An open question is at what point during the mRNA transport cycle KIF1C undergoes LLPS. Given that the purified IDR undergoes LLPS near the estimated KIF1C endogenous concentration, we propose that KIF1C is on the edge of LLPS in cells and its formation of RNA-containing condensates is spatially regulated. That is, we hypothesize that individual KIF1C proteins walk to the plus ends of microtubules in the cell periphery where their accumulation results in a high local concentration that allows LLPS on the microtubule through an IDR-based mechanism (Fig. 9). At concentrations above endogenous levels, whether by overexpression or concentration of the cytoplasm upon hypotonic treatment, KIF1C readily undergoes phase separation, and the resulting condensates display several biophysical characteristics consistent with a liquid-like state including round morphology, fusion and fission, and dynamic exchange of internal components with the surrounding solution.

Several lines of evidence support the proposal that KIF1C LLPS occurs at the end of the transport journey when the motor-cargo complex reaches the plus ends of microtubules (Fig. 9). First, endogenous KIF1C protein is localized diffusely throughout the cell and only a few small droplets are observed in the cell periphery^45,49^. Second, mRNAs whose localization at the cell periphery is dependent on KIF1C can be transported in the form of single copies and then coalesce into larger clusters once they reach their destination in cell protrusions^43,85^. Finally, individual KIF1C proteins are superprocessive and readily reach and accumulate at the plus ends of microtubules^56,57^.

An outstanding question is what effect KIF1C’s LLPS and transport have on the processing, translation, and/or stability of the mRNAs it delivers to the cell periphery. Previous work demonstrated that mRNA in clusters at the tips of retracting cell protrusions are translationally silent^85^, suggesting that KIF1C condensates provide a platform for regulating RNA activity.

Support for this model comes from proximity labeling experiments using BioID fused to the KIF1C C-terminal tail domain^57^. Indeed, nearly half of the proteins identified in the KIF1C- BioID interactome are RBPs, many of which are involved in mRNA decay (Fig S10 E), suggesting an inhibitory role of KIF1C condensates in translational regulation during cell activities such as migration^16,49^.

## Materials and Methods

### Molecular cloning

A plasmid for expression of full-length human KIF1C (pKIF1C-GFP) was a gift from Anne Straube (Addgene plasmid #130977^107^). All truncated versions of KIF1C (see Resource Table), unless otherwise stated, were constructed by PCR (Q5 High-Fidelity DNA Polymerase, NEB cat#M0491S) and ligation or Gibson isothermal assembly (NEBuilder HiFi DNA Assembly Master Mix, cat# E2621S). To obtain a constitutively-active, dimeric KIF1C motor domain [KIF1C(MD)-LZ-mNG], a leucine zipper (LZ) motif was included to ensure dimerization as truncated kinesin-3 motors can form weak dimers, as shown in previous studies^108^. Plasmids for expression of fusion proteins in mammalian cells utilize the cytomegalovirus (CMV) promoter. Fragments for bacterial expression were cloned into pMW96 plasmid (gift of Tarun Kapoor, Addgene plasmid #178066) which uses the T7 promoter to drive protein expression in Rosseta2

*E. coli* cells. All plasmids were verified by DNA sequencing.

### Protein structure prediction

The intrinsically disordered region (IDR) was predicted using IUPred2 (https://iupred2a.elte.hu). The prion-like domain (PLD) was predicted using PLAAC (http://plaac.wi.mit.edu/) with parameters set to core length = 60, α = 100. Results from IUPred2 (ANCHOR score) and PLAAC (HMM.PrD-like score) were visualized in the same plots by custom scripts in R (https://cran.r-project.org).

### Cell culture and transient transfection

COS-7 [male *Ceropithecus aethiops* (African green monkey) kidney fibroblast, RRID: CVCL_0224] cells were grown in Dulbecco’s Modified Eagle Medium (Gibco) supplemented with 10% (vol/vol) Fetal Clone III (HyClone) and 2 mM GlutaMAX (L-alanyl-L-glutamine dipeptide in 0.85% NaCl, Gibco). hTERT-RPE1 cells (female *Homo sapiens* retinal pigment epithelium, RRID: CVCL_4388) were grown in DMEM/F12 with 10% (vol/vol) FBS (HyClone), 0.5 mg/ml hygromycin B, and 2 mM GlutaMAX (Gibco). All cell lines were purchased from American Type Culture Collection and grown at 37 ℃ with 5% (vol/vol) CO_2_. All cell lines are checked annually for mycoplasma contamination and COS-7 cells were authenticated through mass spectrometry (the protein sequences exactly match those in the *Ceropithecus aethiops* genome).

For transfection of plasmid DNA in 6-well plates (2 mL final volume of media each well), 1 ug plasmid DNA and 3 uL TransIT-LT1 transfection reagent (Mirus, Cat#2300) were diluted in 100 uL Opti-MEM Reduced-Serum Medium (Gibco, cat#31985062) to make the transfection mixture. The mixture was added to media in each well immediately after seeding cells (0.5-1x10^5^ cells). 24 hr after transfection, the cells were fixed at 60-70% confluency. The amount of each reagent was scaled up or down when transfecting different volume of cells.

Synthesized RNA oligos (GU-rich oligos: Cy5-mGmUmGmUmGmUmGmUmGmUmGmU; polyA oligos: Cy5-mAmAmAmAmAmAmAmAmAmAmAmAmA) were custom-ordered from MilliporeSigma and were resuspended as 100 μM stock solutions in RNase-free water. The RNA oligos are labeled by Cyanine5 (Cy5) at their 5’ ends and have methylation at each nucleotide to prevent degradation. The cells were transfected by plasmid DNA encoding KIF1C(ST) or KIF1C(IDR) so that KIF1C condensates were already present in cells and then 24 hr later, were transfected with the synthesized RNA oligos (MilliporeSigma). The oligos at various concentrations and 4 uL Lipofectamine RNAiMAX transfection reagent (Invitrogen, Cat# 13778100) were separately diluted in 500 uL Opti-MEM (Gibco, Cat#31985062) and incubated for 5 min at room temperature. Then the two tubes were mixed to make a final mixture of 1 mL and incubated for 15 min at room temperature. 1 mL culture media was removed from each well and replaced with 1 mL transfection mixture. Cells were incubated at 37℃ from various length of time (up to 4 hr) before fixation and further processing.

### Immunofluorescence

For paraformaldehyde fixation, the cells were rinsed with PBS and fixed in 3.7% (vol/vol) paraformaldehyde (ThermoFisher Scientific) in PBS for 10 min at room temperature. Fixed cells were permeabilized in 0.2% Triton X-100 in PBS for 5 min. For methanol-fixed cells, the cells were rinsed with PBS, then fixed and permeabilized at the same time in pre-chilled methanol for 8 min in -20 ℃. After fixation and permeabilization, the cells were blocked with 0.2% fish skin gelatin in PBS for 5 min. Primary antibodies were applied in 0.2% fish skin gelatin in PBS overnight at 4 ℃ in a custom humidified chamber. Following 3X washes with PBS with 0.2% fish skin gelatin, secondary antibodies were applied in 0.2% fish skin gelatin in PBS for 1 hr at room temperature in the dark. Nuclei were stained with 10.9 mM 40,6-diamidino-2-phenylindole (DAPI) and the coverslips were mounted using Prolong Gold (Invitrogen). Images were acquired on an inverted epifluorescence microscope (Nikon TE2000E) with a 40x, 0.75 NA, a 60x, 1.40 NA oil-immersion, or a 100x, 1.40 NA objective and a CoolSnap HQ camera (Photometrics).

### Live-cell imaging

For the cytoplasm dilution assay, cells were seeded in 35 mm glass-bottom dishes (Matek, Cat# P35G-1.5-14-C). 16-24 hr post-transfection, the cells were washed with and then incubated in Leibovitz’s L-15 medium (Gibco) and imaged at 37 °C in a temperature-controlled and humidified stage-top chamber (Tokai Hit) on a Nikon X1 Yokogawa Spinning Disk Confocal microscope with a 60x, 1.49 NA oil-immersion objective, and a Andor DU-888 camera. Image acquisition was controlled with Elements software (Nikon). To minimize stage drifting during media exchange, a syringe was attached to the microscope stage using modeling clay to keep it static relative to the 35 mm glass-bottom dish. The syringe was used to remove media from the dish, and then new media was added by gentle pipetting. During live-cell imaging, cells were imaged in L-15 medium for more than 1 min to let the imaging process stabilize, washed once with hypotonic medium (Leibovitz’s L-15 medium diluted by 1:4 with sterile water), and then incubated in the hypotonic medium to observe the change of LLPS. The dish was then washed once with the isotonic medium (Leibovitz’s L-15 medium), and incubated in the isotonic medium to observe recovery.

For fluorescence recovery after photobleaching (FRAP) experiments, cells in glass-bottom dishes (Matek, Cat# P35G-1.5-14-C) were observed on a Nikon A1R line scan confocal microscope (Biomedical Research Core Facilities, University of Michigan). Prebleach images were acquired, followed by photobleaching at 100% laser power at 488 nm and 80% laser power at 561 nm for 1 sec, and then fluorescence recovery images were collected over time (1 frame every 3 sec). Afterwards, the images were corrected for photobleaching in Fiji (https://imagej.net/software/fiji/). Fluorescence intensity in each frame was quantified in Fiji and analyzed using R scripts. The half-time of fluorescence recovery was calculated by fitting the recovery curve after the point of bleaching to an exponential curve:

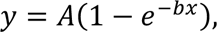

where y is normalized intensity, x is time after bleaching. Once parameters A and b were calculated in R, the time constant was defined as τ_1/2_ = 0.693/b. The mobile fraction was calculated by averaging the normalized intensity after the point where the recovery reaches its plateau.

For microinjection of RNase A, COS-7 cells expressing KIF1C(ST)-mNG plasmid were washed with Leibovitz’s L-15 medium and then incubated in Leibovitz’s L-15 medium. The dish was mounted on the stage of Nikon Eclipse (TE2000-U) microscope and images were obtained with a CoolSNAP ES2 camera at room temperature. The microinjection was performed with glass capillaries (Eppendrof Femtotips, Cat# 930000035) loaded with 5 mg/mL of RNase A (Roche, Cat# 10109142001) mixed with Dextran-TAMRA in PBS. Before mixing, the RNase and Dextran-TAMRA aliquots were spun at 4 ℃ for 10 min to remove any precipitates. For control experiments, only Dextran-TAMRA in PBS was loaded in glass capillaries. The micromanipulator (Eppendorf InjectMan NI 2) was positioned and microinjection was performed using microinjector (Eppendorf FemtoJet) with a 40x dry objective lens to facilitate immediate visualization and image acquisition before and after microinjection. 40 hPa of compensation pressure (Pc) and 95 hPa injection pressure (Pi) for 1.0 second injection time (Ti) was used for each microinjection.

### Protein purification from bacteria and evaluation of liquid-liquid phase separation (LLPS) in vitro

KIF1C fragments tagged with 6xHis, SUMO and mNG in the pMW96 vector were transformed into Rosetta2 cells. Single colonies were cultured overnight, reinoculated the next day to grow to log phase, and protein expression was induced by adding 1 mM isopropyl β-D-thiogalactoside (IPTG) (Invitrogen, Cat# 15529019). After 16 hr, the cells were collected by centrifugation at 20,000 x g for 3 min. Bacterial pellets were resuspended in high-salt lysis buffer (500 mM NaCl,

1% TritonX-100, 50 mM Tris pH8.0, 10 mM MgCl_2_, 0.1 mg/mL lysozyme, 1x Benzonase nuclease) by thorough vortexing. After incubation at 37℃ for 30 min, the mixture was centrifuged at 20,000 x g for 15 min at 4℃. The soluble fraction (supernatant) was collected and mixed with Ni-NTA resin at 4℃ overnight with slow rotation. The protein-bound resin was washed 3X with wash buffer (500 mM NaCl, 1% TritonX-100, 50 mM Tris pH8.0, 10 mM MgCl_2_, 25 mM Imidazole, 1 mM DTT) and 6xHis-SUMO-TEV-KIF1C-mNG protein was eluted by elution buffer (500 mM NaCl, 1% TritonX-100, 50 mM Tris pH8.0, 10 mM MgCl_2_, 250 mM Imidazole, 1 mM DTT). Imidazole was removed by 3 cycles of concentration and dilution using Vivaspin (Millipore Sigma, Cat# GE28-9322-25). TEV enzyme was added at about 1/10 of the target protein mass and incubated overnight. The reaction mix was then incubated with Ni-NTA resin to retain the cleaved His-SUMO tag. KIF1C-mNG protein was collected in the flow- through and concentrated with Vivaspin. The protein solution was brought to 10% glycerol, aliquoted, and snap frozen in -80 ℃ freezer. The purity and concentration of the purified proteins were analyzed by SDS-PAGE and SimpleBlue SafeStain (Invitrogen, Cat#LC6060).

Purified protein in high salt buffer (500 mM NaCl, 1% TritonX-100, 50 mM Tris pH8.0, 10 mM MgCl_2_, 1 mM DTT) was mixed with low-salt buffer (0 mM NaCl, 1% TritonX-100, 50 mM Tris pH8.0, 10 mM MgCl_2_, 1 mM DTT) to reduce NaCl concentration to desired level. The solution was introduced into a flow chamber (10 μL volume) assembled by attaching a clean #1.5 coverslip (Thermo Fisher Scientific) to a glass slide (Thermo Fisher Scientific) with two stripes of double-sided tape. The flow chamber was first rinsed using a solution with NaCl concentration equivalent to the protein solution. Images were acquired on an inverted epifluorescence microscope (Nikon TE2000E) with a 100x, 1.40 NA objective and a CoolSnap HQ camera (Photometrics) at room temperature.

### Generation of KIF1C-KO cells by CRISPR/Cas9-mediated genome editing

Knockout of KIF1C from the genome of hTERT-RPE1 cells was performed as described previously^109^. The 20-nt sgRNA sequences were designed by the CRISPR guide RNA design tool in Benchling (https://benchling.com). The oligo pairs [forward strand: 5’- aaacCGGTGAAAGTGGCAGTGAGGc-3’; reverse strand: 5’- caccgCCTCACTGCCACTTTCACCG-3’] were synthesized (IDT), annealed, and ligated into plasmid eSpCas9(1.1). The product plasmid was transfected into wild type hTERT-RPE1 cells using TransIT-LT1 transfection reagent. 48 hours post-transfection, the cells were selected in 800 μg/mL Geneticin (Gibco) until cells in the untransfected control group died. Single cells were then isolated through flow cytometry (non-fluorescent) and sorted into 96-well plates.

Colonies were expanded and genomic DNA was extracted. The KIF1C sequence close to the sgRNA target site was amplified by PCR and sequenced. KIF1C-KO cell lines identified by DNA sequencing were verified for loss of KIF1C expression by Western Blot using Rb anti- KIF1C (Abcam) antibody.

### Estimation of endogenous KIF1C concentration by Western Blot

hTERT-RPE1 parental cells were trypsinized and harvested by centrifugation at 5000 x g at 4 ℃ for 5 min, washed with cold 1X PBS, and resuspended in cold lysis buffer [25 mM HEPES/KOH, 115 mM potassium acetate, 5 mM sodium acetate, 5 mM MgCl_2_, 0.5 mM EGTA, and 1% (vol/vol) Triton X-100, pH 7.4] with 1 mM ATP, 1 mM phenylmethylsulfonyl fluoride (PMSF), and 1% (vol/vol) protease inhibitor cocktail (P8340, Sigma-Aldrich). Lysates were clarified by centrifugation at 20,000 x g at 4 ℃ for 10 min and the supernatants were mixed with 5X Laemmli buffer and 1: 50 1 mM DTT. The samples were boiled at 95 ℃ for 5 min, cooled to room temperature and briefly centrifuged before loading into SDS-PAGE gels and immunoblotted using Rb anti-KIF1C antibody (Abcam, Cat#ab72238). Purified KIF1C(IDR)- mNG was used to generate a protein standard. The mass of KIF1C was calculated by fitting the intensity of protein band to the KIF1C(IDR)-mNG standard curve. The volume of cytoplasm was estimated by multiplying the number of cells loaded and the average volume of each cell. For example, 1.35 x 10^6^ hTERT-RPE1 cells x 2416 ± 263 μm^3^/cell [mean ± STD, ^110^] = 3.3 μL of cytoplasm loaded into each lane. The endogenous KIF1C concentration was calculated by dividing the mass of KIF1C by the cytoplasmic volume and protein molecular weight, and was averaged across 3 experiments. Note that due to loss of protein during the process of making lysates, the measured value is likely to be an underestimate.

### Single-molecule fluorescence *in situ* hybridization (smFISH)

Single-molecule fluorescence *in situ* hybridization and the sequences of probes against *Rab13* mRNA were based on previous literature^43,111,112^. The primary and secondary probes were annealed and diluted in hybridization buffer (10% Formamide, 1 mg/mL *E.coli* tRNA, 10% Dextran Sulfate, 0.2 mg/mL BSA, 2x Saline-sodium citrate buffer, 2 mM Vanadyl ribonucleoside complex). Cells were washed 3X with PBSM (PBS with 5 mM MgCl_2_), fixed with 3.7% (vol/vol) paraformaldehyde in PBSM for 10 min at room temperature, washed 2X with PBSM, and permeabilized with permeabilization buffer (0.1% TritonX-100 in PBSM) for 10 min. Cells were then washed 2X and incubated with pre-hybridization buffer (10% Formamide in 2x SSC) for 30 min at RT, and the incubated with annealed probes in hybridization buffer overnight at 37 ℃ in a custom-made humidified chamber. The next day, the cells were washed twice with pre-hybridization buffer at 37 ℃ in humidity chamber, and three times with 2x SSC at room temperature. 0.5 μg/mL DAPI staining solution was applied for 1 min at room temperature. The cells were washed three times and finally mounted with ProLong Gold (Invitrogen). Images were acquired on an inverted epifluorescence microscope (Nikon TE2000E) with a 40x, 0.75 NA, a 60x,1.40 NA oil-immersion, or a 100x, 1.40 NA objective and a CoolSnap HQ camera (Photometrics).

### Image analysis

Image analysis was carried out using Fiji/ImageJ (https://fiji.sc). For quantification of KIF1C and KIF16B puncta sizes (Fig. 1 C), puncta that were aggregated in cell protrusions were excluded from quantification as they could not be individually defined.

For analysis of KIF1C enrichment in puncta (Fig. 2), measurements were made on a single cell basis of the mean fluorescence intensity value of all puncta (I_puncta_) and compared to the mean fluorescence intensity of an area adjacent to the puncta (I_diffusive_). The background intensity (I_background_) was measured from neighboring untransfected cells and the final ratio was calculated by the following equation:

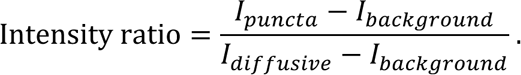

The calculated intensity ratios were plotted as a violin plot in R where each point represents measurement from one cell.

Colocalization between KIF1C puncta and *Rab13* mRNA (Fig. 6 D) was measured by drawing a segmented line across KIF1C puncta and counting the number of *Rab13* mRNA puncta that colocalize with KIF1C puncta along the line. A KIF1C punctum was defined as a fluorescent peak along the line where the intensity value is at least 1.5-fold the average intensity along the line. A colocalizing *Rab13* mRNA peaks was defined as a fluorescence peak within 0.4 μm distance (∼ 3 pixels) of the KIF1C punctum.

When quantifying puncta size of different IDR variants of KIF1C in each cell (Fig. 7 C), a weighted mean was calculated by:

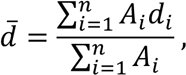

where d_i_ is the diameter of individual puncta in a cell, A_i_ is the corresponding area of puncta. When there are a couple of dominating puncta in a cell that contain most of dense phase KIF1C, this weighted mean (shown above) can represent this situation better than an ordinary arithmetic mean.

## Key Resources Table

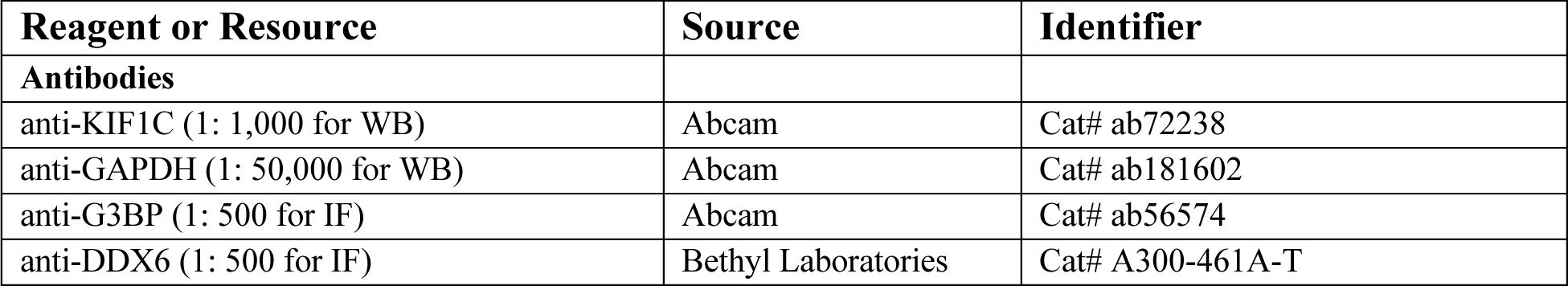
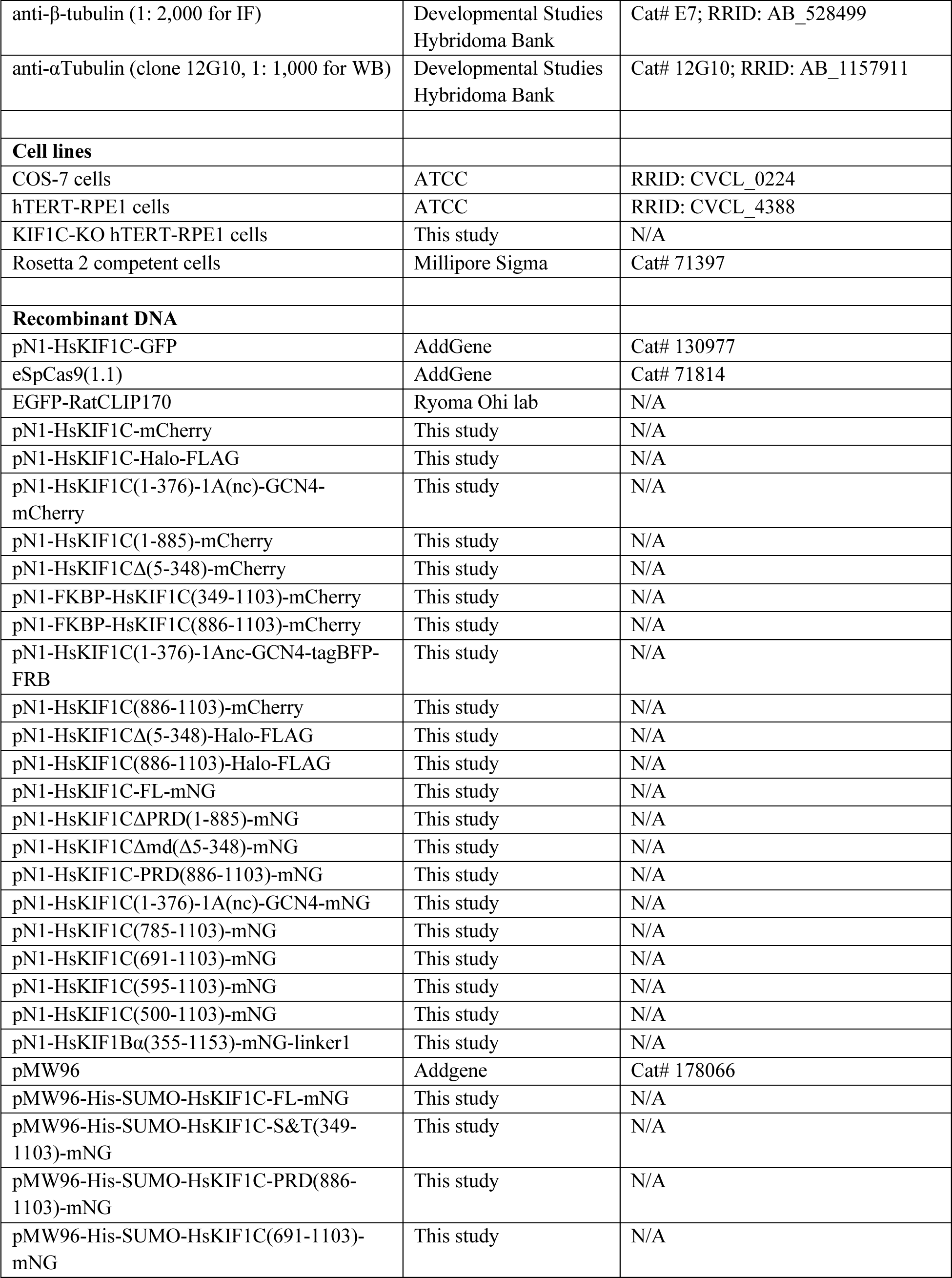
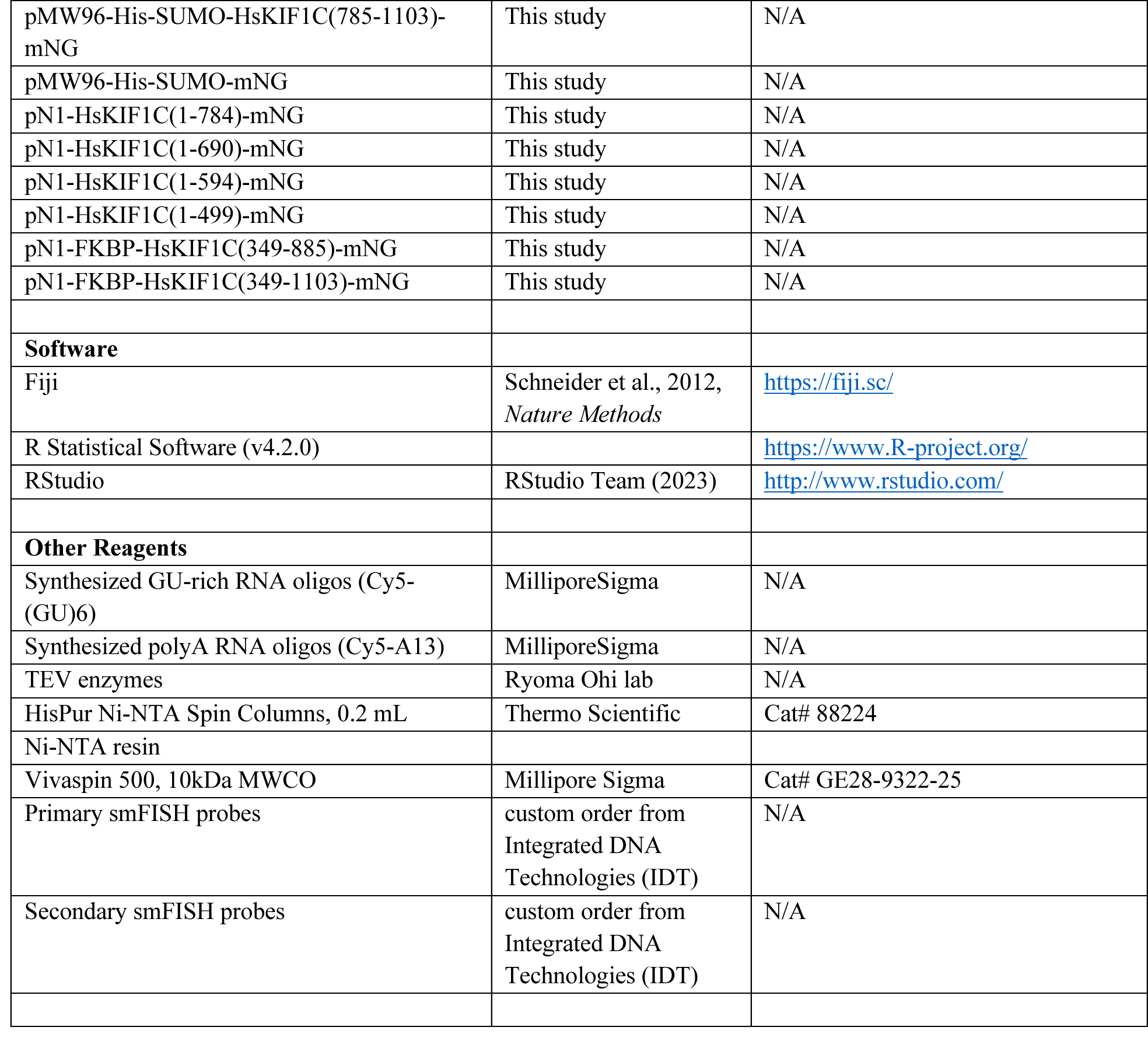

## Acknowledgements

We thank the members of the Verhey, Cianfrocco, DeSantis, Ohi and Sept lab for ideas, discussion, feedback, and support. We thank Zesen Lin and Rami Khoriaty for help with CRIPSR-Cas9 protocols for generating KIF1C-KO cells, Amanda Erwin and Shyamal Mosalaganti for help with experiments using fluorescently-labelled synthetic RNA oligos, and Ye Yuan and Swathi Yadlapalli for help with smFISH. We are grateful to Eric Rentchler in the University of Michigan Biomedical Research Core facilities for training and guidance. This work was supported by funding from the National Institutes of Health to KJV (R35GM131744).

---

**Figure S1.**
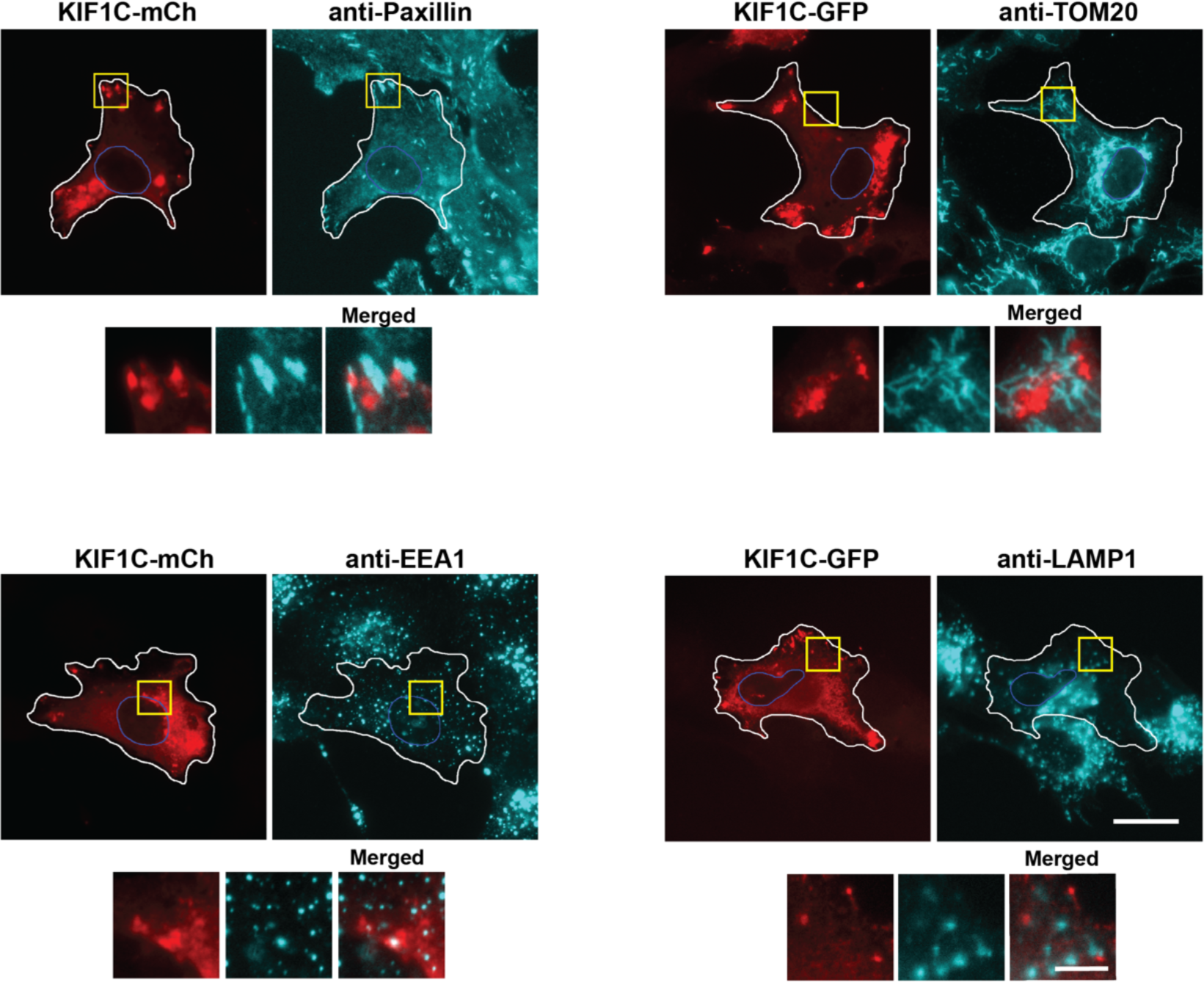
KIF1C puncta do not colocalize with organelle markers. Immunofluorescence of markers for focal adhesions (Paxillin), early endosomes (EEA1), mitochondria (TOM20), and lysosomes (LAMP1) in hTERT-RPE1 cells expressing fluorescently-tagged KIF1C. Representative images are shown. White lines indicate cell boundaries. Blue lines indicate nuclear boundaries. Yellow boxes indicate regions shown in magnified images below. Scale bar: 20 μm for whole cell views, 5 μm for magnified images.

**Figure S2.**
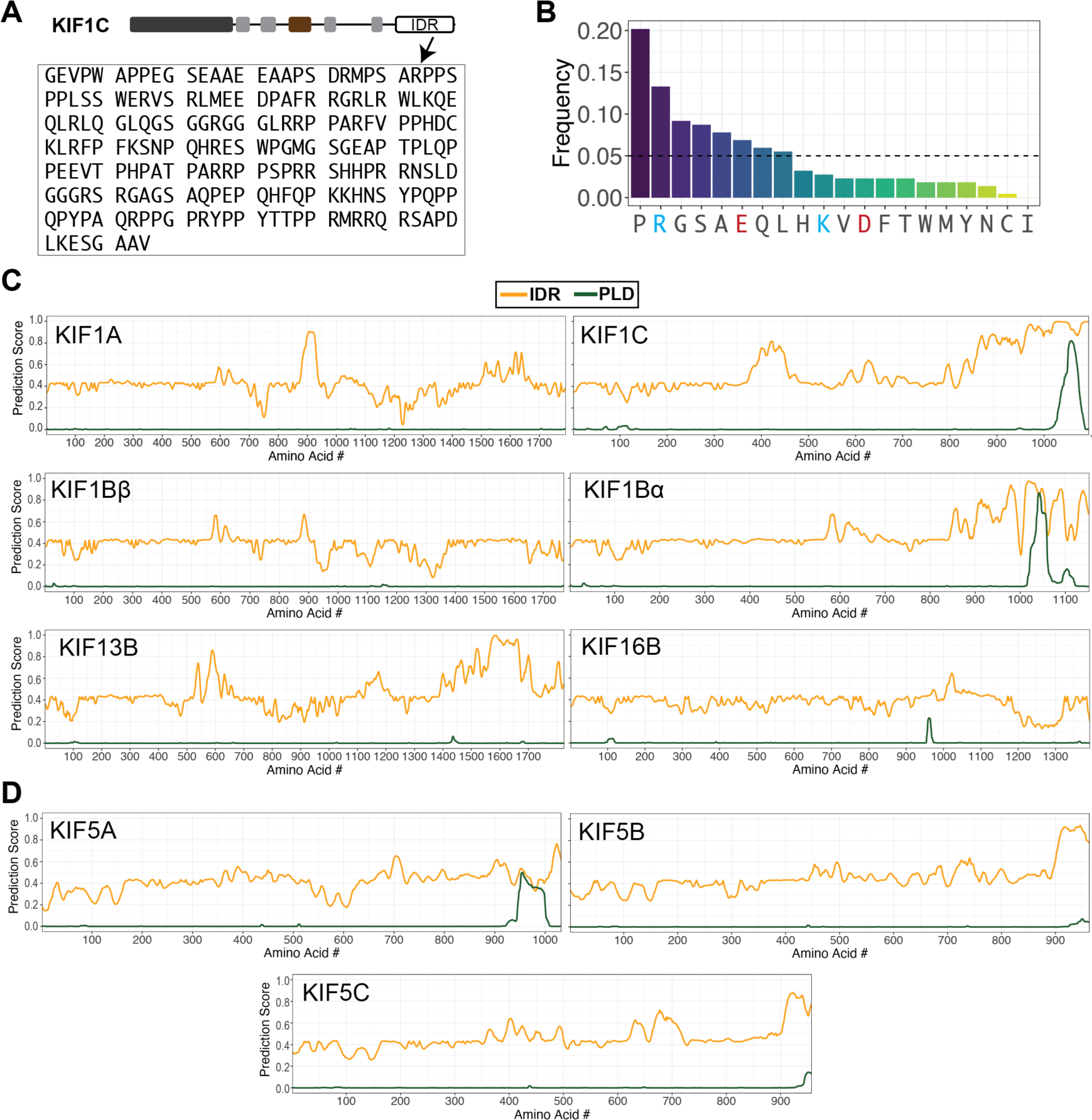
**The KIF1C tail domain is an IDR.** (A) Amino acid sequence of the KIF1C IDR. (B) Frequency of amino acid residues in the KIF1C IDR. The horizontal dashed line indicates the frequency of 0.05. For the x-axis, the positively charged residues R and K are labelled blue; the negatively charged residues E and D are labelled red. (C,D) IUPred and PLAAC predictions of IDR and PLD, respectively, for (C) kinesin-3 family members and (D) kinesin-1 family members. x-axis: amino acid residue number; y-axis: predicted probability of the given residue being part of an IDR (orange line) or a PLD (green line).

**Figure S3.**
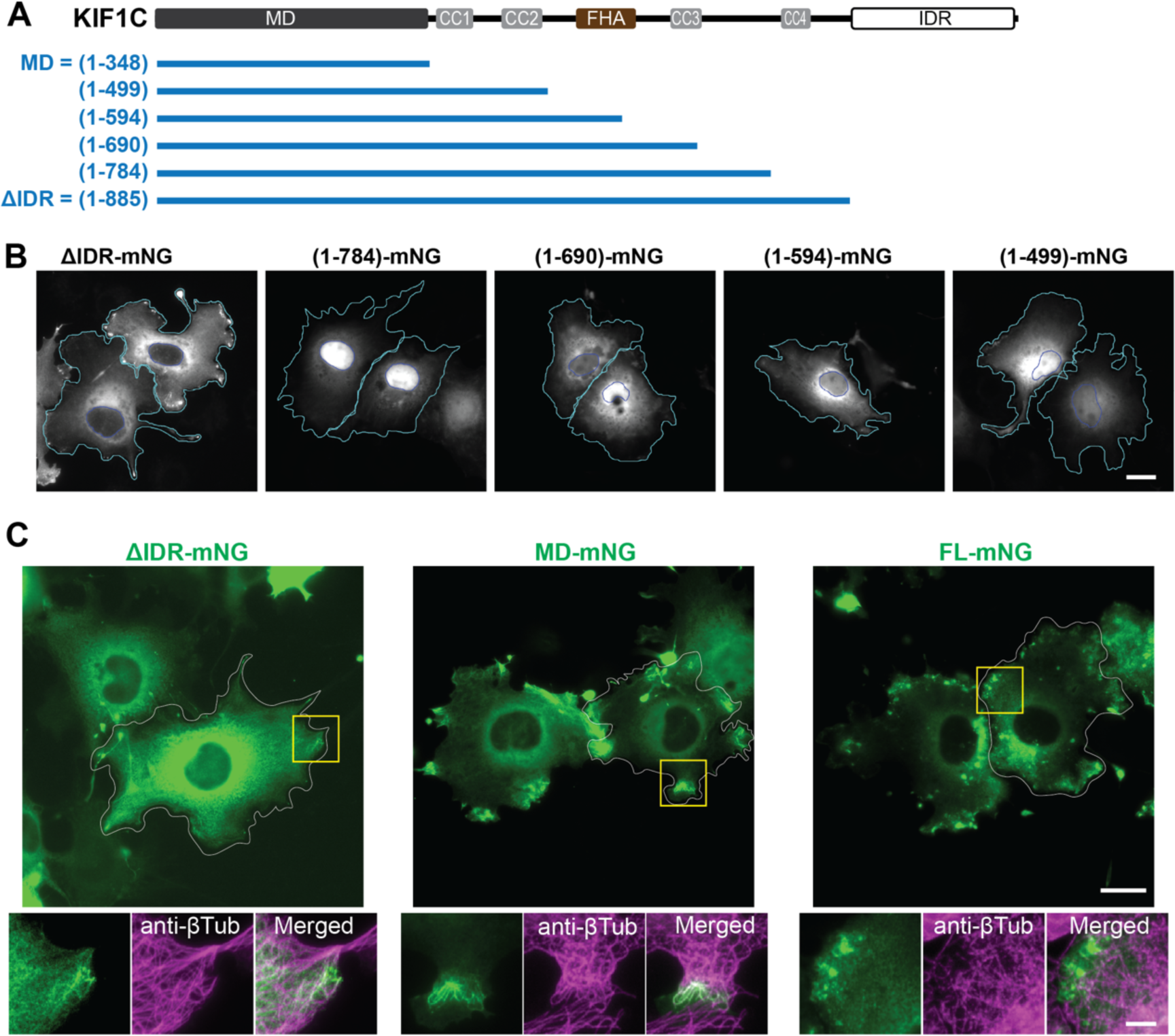
**The IDR is required for KIF1C puncta formation in cells.** (A) Schematic of KIF1C serial truncations from the C-terminus. (B) Representative images of KIF1C truncations in COS-7 cells. Scale bar: 20 μm. Cyan lines indicate cell boundaries. Blue lines indicate nuclear boundaries. (C) Immunofluorescence of microtubules (anti-βTubulin) in COS-7 cells expressing KIF1C(ΔIDR)-mNG, KIF1C(MD)-mNG, or KIF1C(FL)-mNG. Scale bar: 20 μm for whole cell views, 5 μm for magnified images.

**Figure S4.**
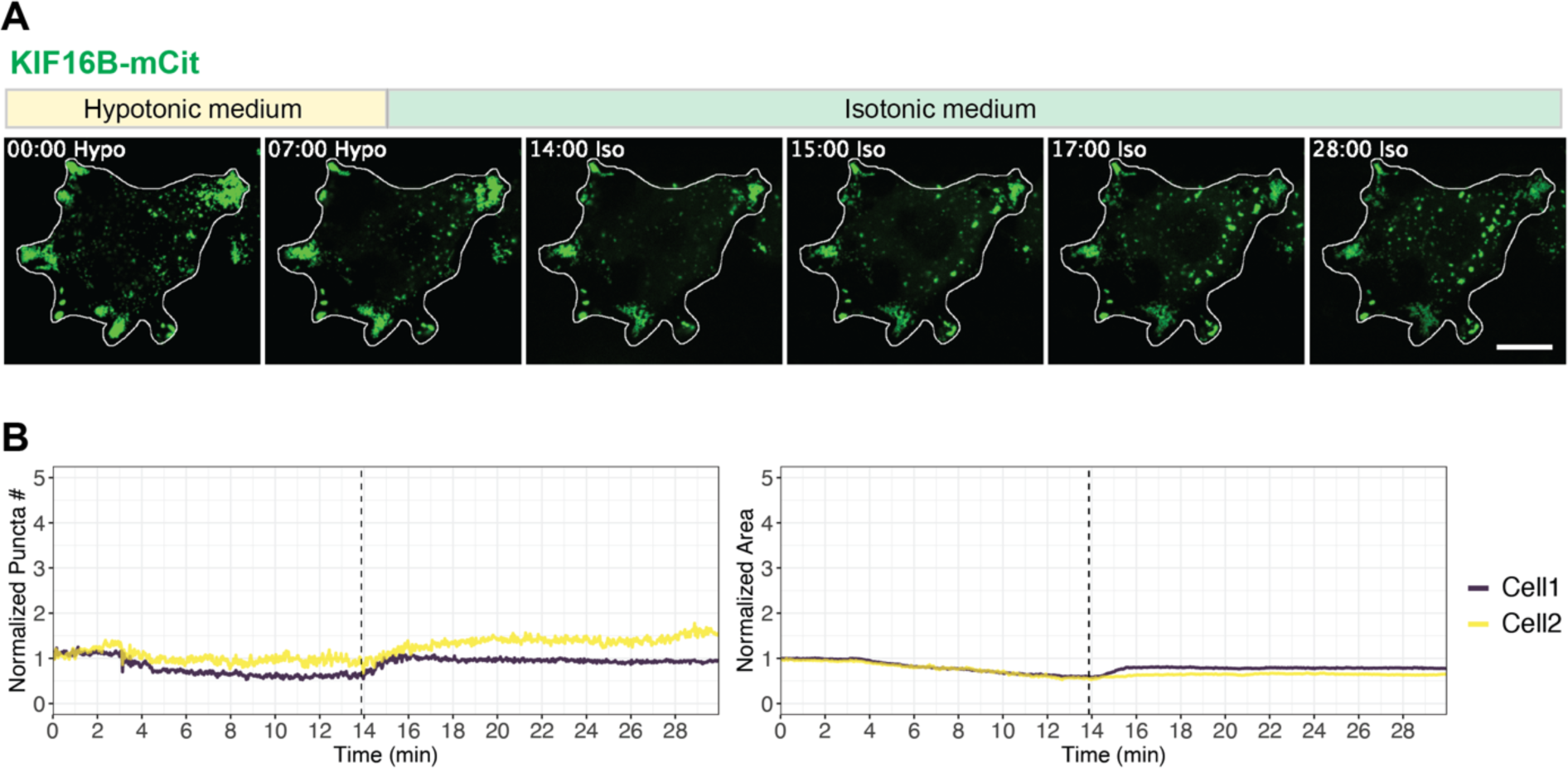
**KIF16B does not change during the cytoplasm dilution assay.** (A) Representative images of mCit-tagged KIF16B localization before treatment, during hypotonic treatment, and upon return to isotonic media. Scale bar: 20 μm. Time label is [min:sec]. (B) Quantification of change of puncta number (left) and total area of puncta (right) over time in the cytoplasm dilution assay. x axis: time, with a vertical dashed line indicating the time point of switching from the hypotonic media to the isotonic media. y axis: puncta number or puncta area normalized against the initial state in the first frame of live-cell imaging. The example cell in (A) is Cell 1 in the plots.

**Figure S5.**
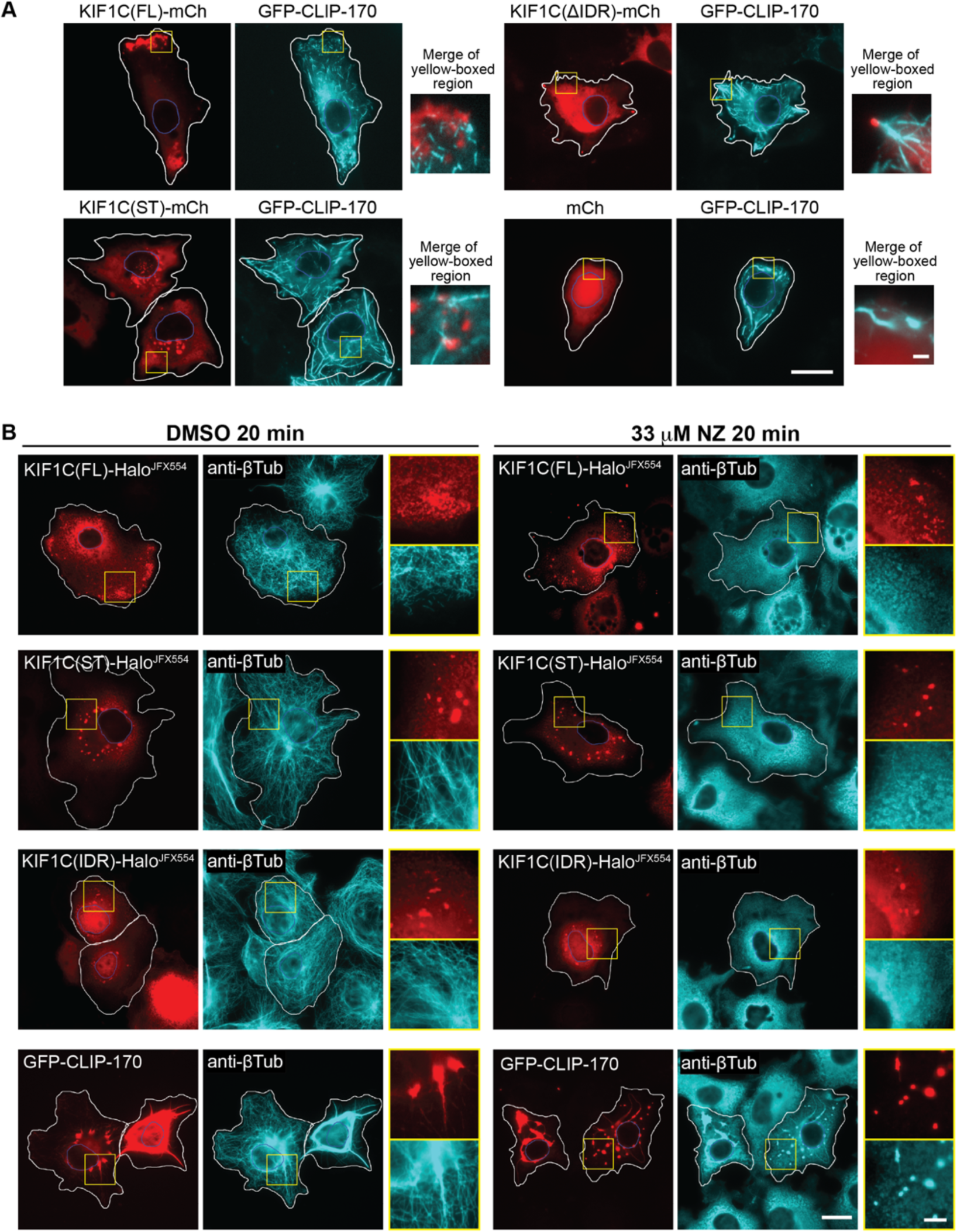
**KIF1C condensates do not colocalize with CLIP-170 or tubulin.** (A) Localization of mCh (control) or mCh-tagged KIF1C constructs (FL, ST, and ΔIDR) co-expressed with GFP-tagged CLIP-170 in hTERT-RPE1 cells. Representative images are shown. White lines indicate cell boundaries. Blue lines indicate nuclei boundaries. Yellow boxes indicate the regions displayed in the magnified images to the right. Scale bars: 20 μm for whole cell images, 2 μm for magnified images. (B) Immunofluorescence of anti-βTubulin in COS-7 cells expressing Halo^JFX554^–tagged KIF1C constructs (FL, ST, and IDR) or GFP-tagged CLIP-170. Cells were treated with (left) DMSO or (right) 33 μM nocodazole (NZ) for 20 min. White lines indicate cell boundaries. Yellow boxes indicate the regions displayed in the magnified images to the right. Scale bars: 20 μm for whole cell images, 5 μm for magnified images.

**Figure S6.**
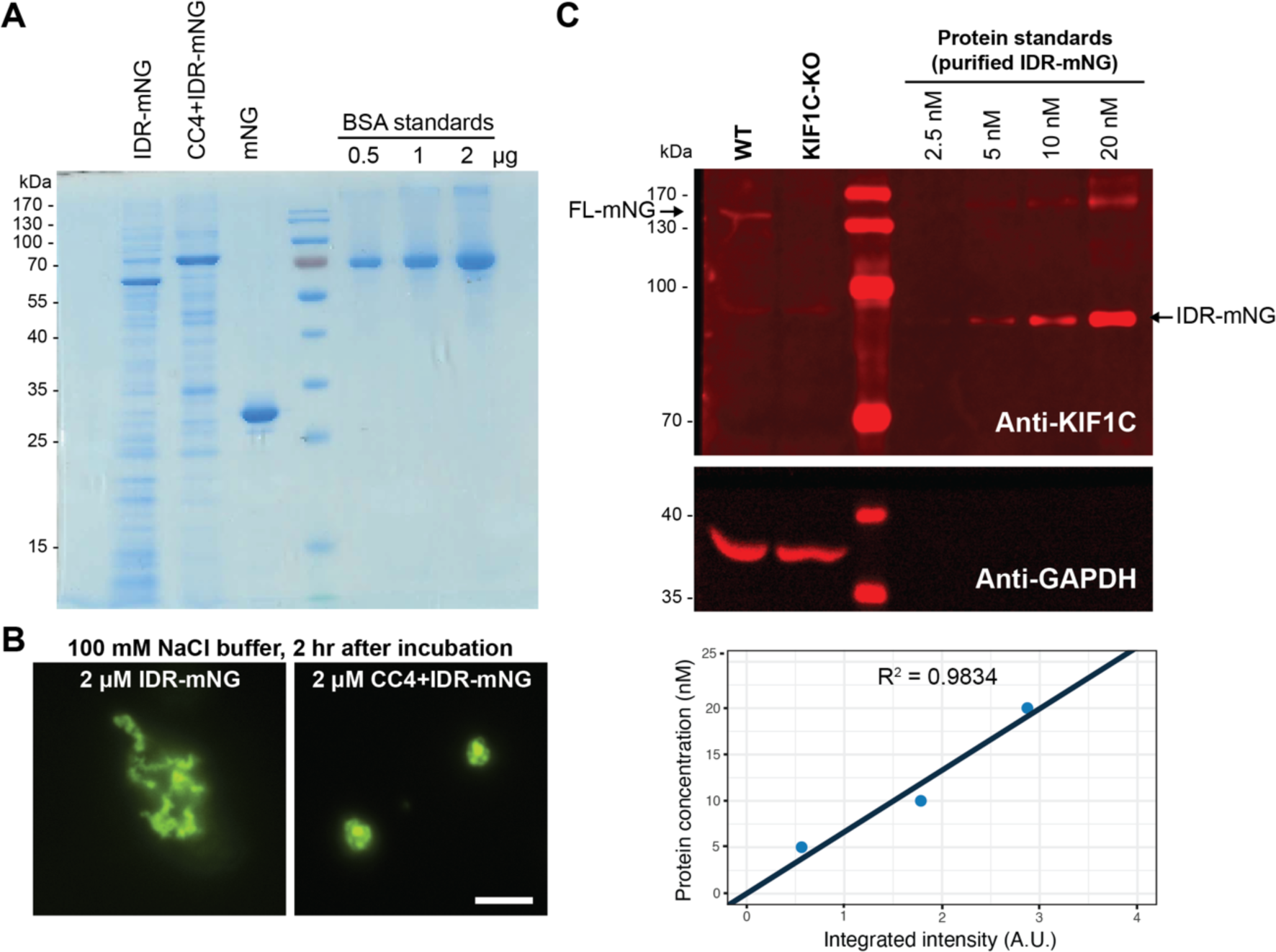
**LLPS of purified IDR and CC4+IDR proteins.** (A) Purified KIF1C(IDR)-mNG, KIF1C(CC4+IDR)-mNG, and mNG separated by SDS-Page. Bovine serum albumin (BSA) was run as protein standards. (B) Representative images of purified KIF1C(IDR)-mNG and purified KIF1C(CC4+IDR)-mNG imaged after prolonged incubation (2 hours). Scale bar: 5 μm. (C) Estimation of endogenous KIF1C concentration in hTERT-RPE cells by western blot. Cell lysates of WT and KIF1C-KO hTERT-RPE1 cells were probed by western blotting with an antibody against KIF1C and against GAPDH (loading control). Purified KIF1C(IDR)-mNG was used as protein standards to generate a standard curve (bottom) for calculating the concentration of endogenous KIF1C.

**Figure S7.**
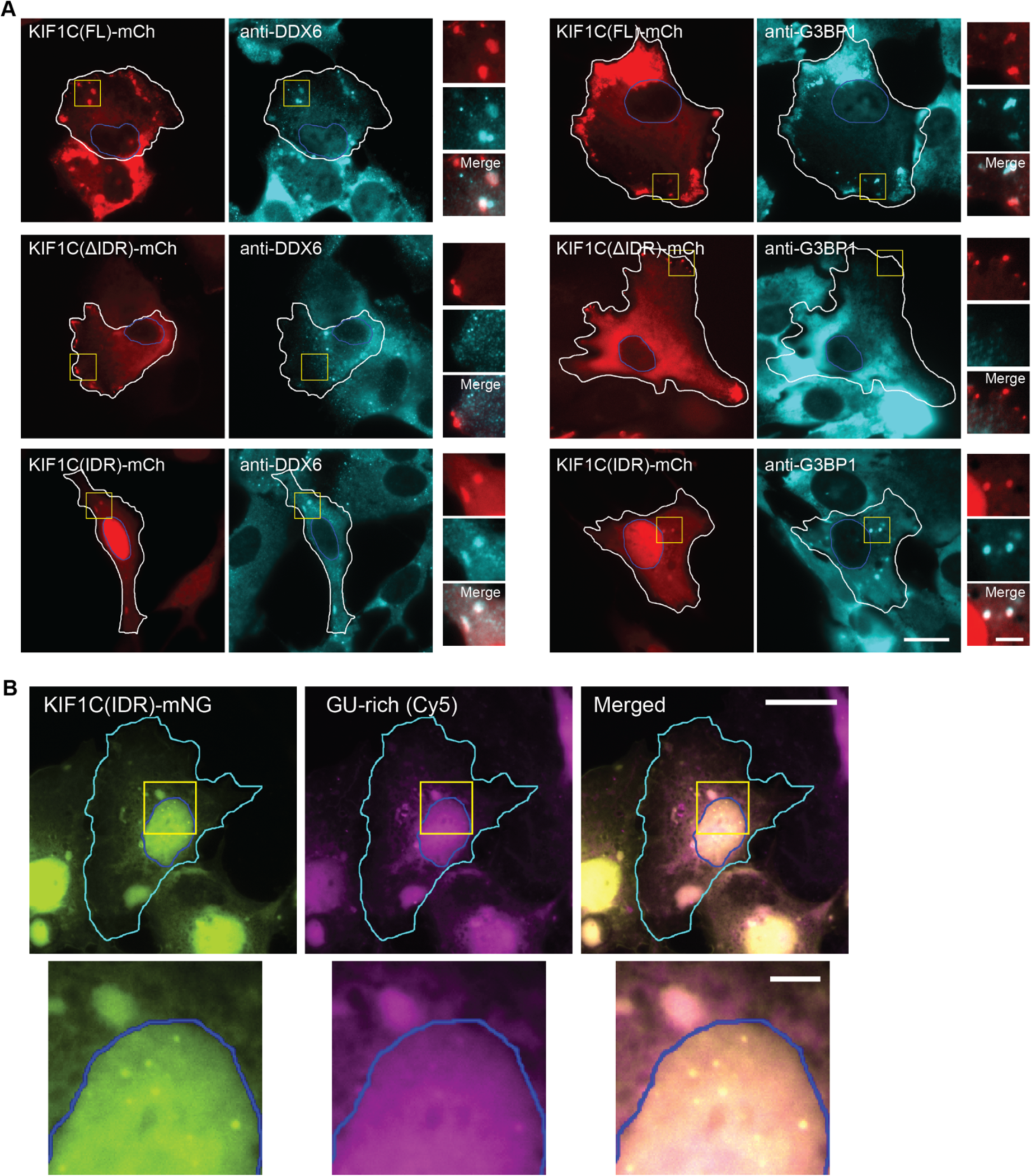
**KIF1C colocalizes with endogenous RNA granules and exogenous RNA.** (A) Immunofluorescence for markers for P-bodies (anti-DDX6) and stress granules (anti-G3BP1) in hTERT-RPE1 cells expressing mCh-tagged KIF1C(FL), KIF1C(ΔIDR), or KIF1C(IDR). Representative images are shown. White lines indicate cell boundaries. Blue lines indicate nuclear boundaries. Yellow boxes indicate the regions displayed in the magnified images to the right. Scale bar: 20 μm for whole cell views, 5 μm for magnified images. (B) Representative images of Cy5-labelled GU-rich RNA oligos introduced into COS-7 cells expressing KIF1C(IDR)-mNG. Yellow boxes indicate the regions displayed in the magnified images on the bottom. Cyan lines indicate cell boundaries. Blue lines indicate nuclear boundaries. Scale bars: 20 μm for whole cell views, 5 μm for magnified images.

**Figure S8.**
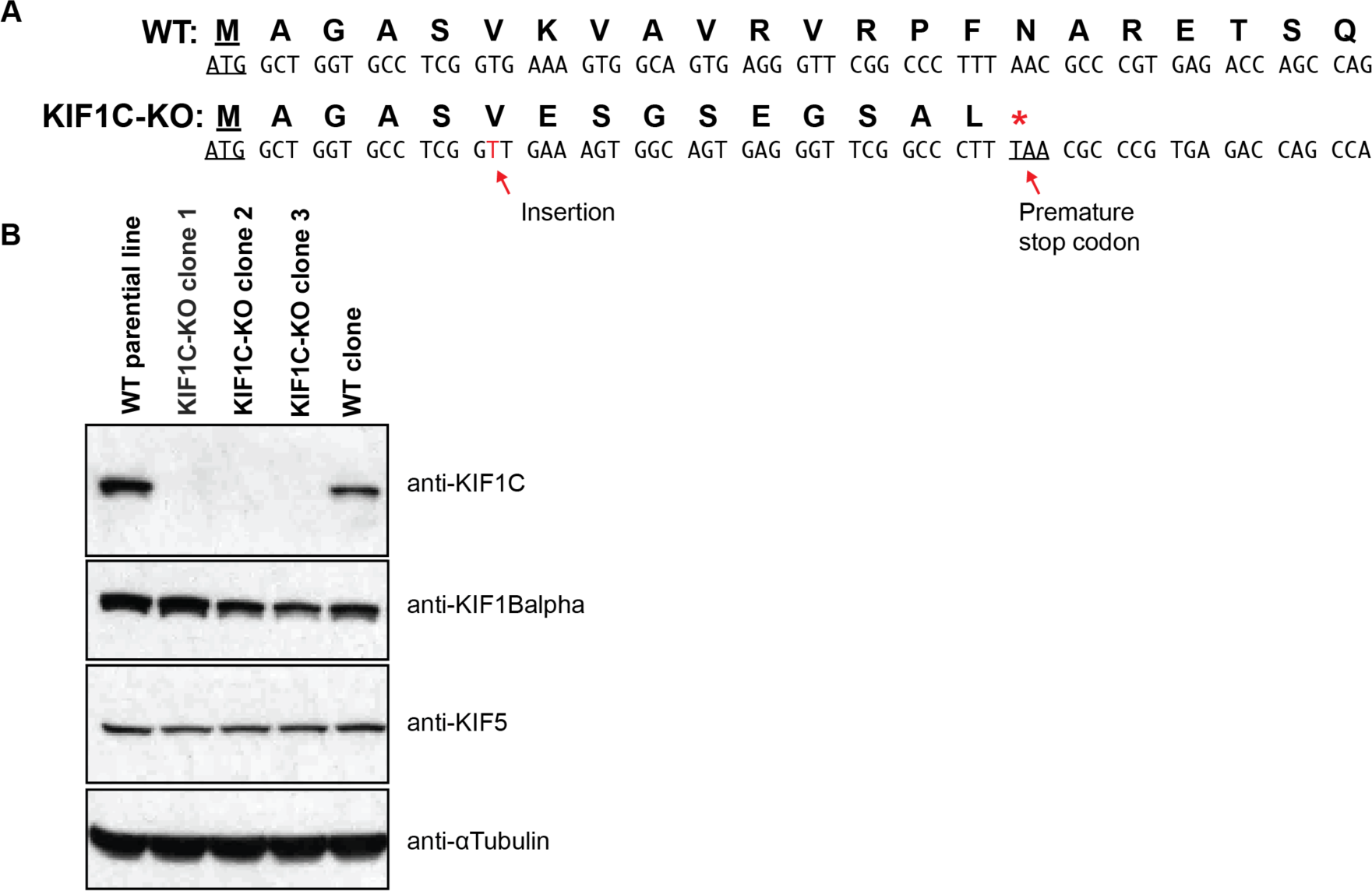
**Verification of KIF1C-KO in hTERT-RPE1 cells.** (A) DNA and protein sequences from the start codon (underlined) in WT (top) and KIF1C-KO (bottom) hTERT- RPE1 cells. The same insertion of T (red text) was found in all 3 clones of KIF1C-KO cells, which results a premature stop codon (underlined and with a red asterisk). (B) Western blot showing the absence of KIF1C protein in all three KIF1C-KO clones.

**Figure S9.**
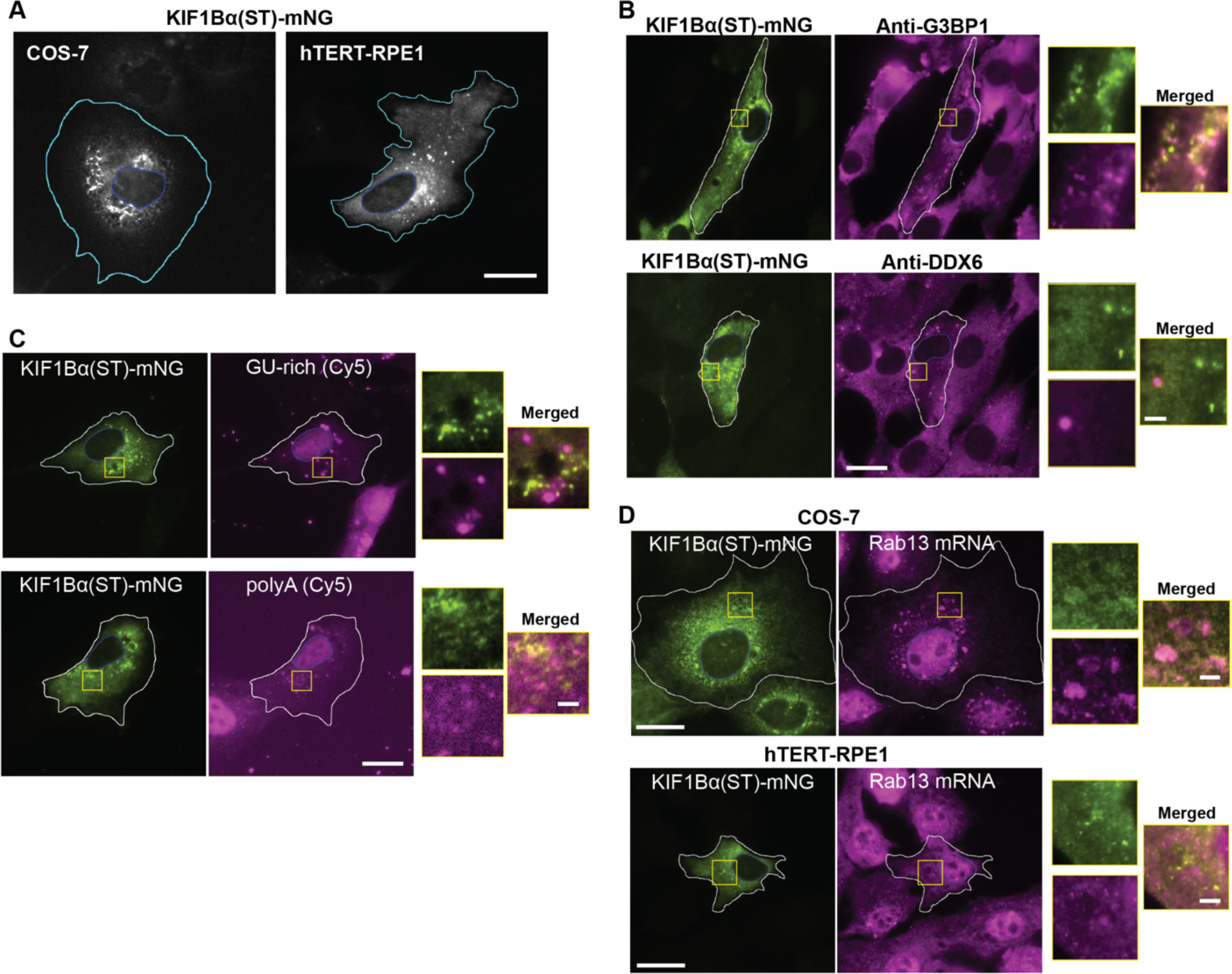
**Characterization of KIF1Bα(ST) puncta.** (A) Localization of KIF1Bα(ST)-mNG in COS-7 cells and hTERT-RPE1 cells. Representative images are shown. Cyan lines indicate cell boundaries. Scale bar: 20 μm. (B) Immunofluorescence for markers for stress granules (anti-G3BP1) or P-bodies (anti-DDX6) in hTERT-RPE1 cells expressing KIF1Bα(ST)-mNG. Representative images are shown. White lines indicate cell boundaries. Yellow boxes indicate the regions displayed in the magnified images to the right. Scale bar: 20 μm for whole cell views, 5 μm for magnified images. (C) Representative images of Cy-5-labelled GU-rich or polyA RNA oligos introduced into hTERT-RPE1 cells expressing KIF1Bα(ST)-mNG. White lines indicate cell boundaries. Yellow boxes indicate the regions displayed in the magnified images to the right. Scale bars: 20 μm for whole cell views, 2 μm for magnified images. (D) Representative images of KIF1Bα(ST)-mNG localization in COS-7 cells (top) or hTERT-RPE1 cells (bottom) with smFISH for endogenous *Rab13* mRNA. White lines indicate cell boundaries. Blue lines indicate nuclear boundaries. Yellow boxes indicate the regions displayed in the magnified images to the right. Scale bars: 20 μm for whole cell views, 2 μm for magnified images.

**Figure S10.**
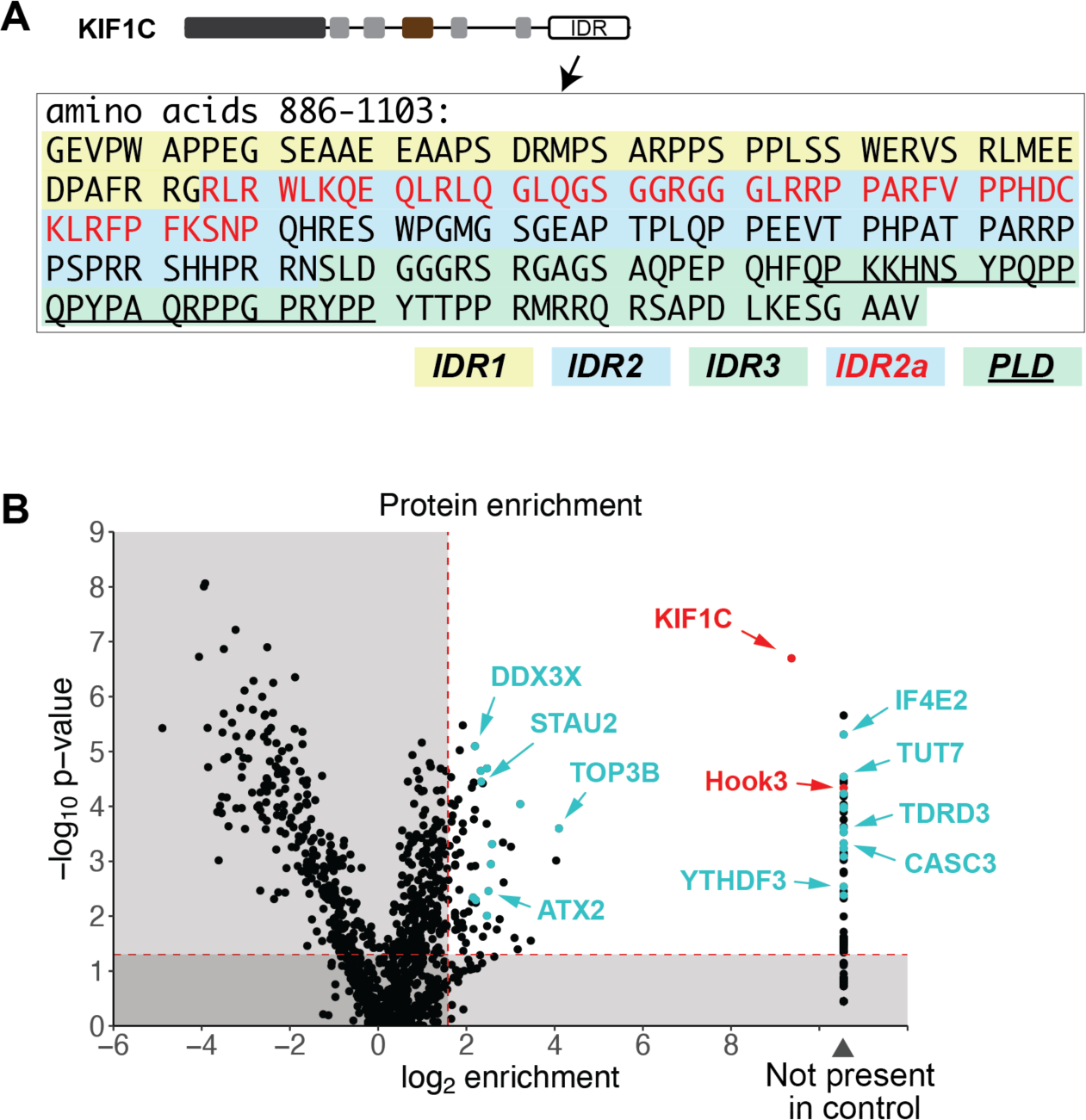
**KIF1C-BioID identified numerous RBDs as interaction partners.** (A) Amino acid sequence of the KIF1C IDR with IDR1, IDR2, and IDR3 subregions color-coded. Subregion IDR2a within IDR2 is highlighted by red text. Subregion PLD within IDR3 is underlined. (B) Volcano plot showing proteins identified in KIF1C-BioID experiments. Image modified from Kendrick et al., 2019, Figure 1D (reference 57). Proteins with an enrichment ratio > 3 and a p-value < 0.01 are included in the list. The interaction between KIF1C and Hook3 proteins (red text) has been characterized by Siddiqui et al., 2019 and Kendrick et al., 2019 (references 56 and 57). The cyan spots indicated RBPs identified in the KIF1C interactome. The cyan text indicates RBPs involved in mRNA processing or decay.

